# Functional hyperemia drives fluid exchange in the paravascular space

**DOI:** 10.1101/838813

**Authors:** Ravi Teja Kedarasetti, Kevin L. Turner, Christina Echagarruga, Bruce G. Gluckman, Patrick J. Drew, Francesco Costanzo

## Abstract

Maintaining the ionic and chemical composition of the extracellular spaces in the brain is extremely important for its health and function. However, the brain lacks a conventional lymphatic system to remove metabolic waste. It has been proposed that the fluid movement through the paravascular space (PVS) surrounding penetrating arteries can help remove metabolites from the brain. The dynamics of fluid movement in the PVS and its interaction with arterial dilation and brain mechanics are not well understood. Here, we performed simulations to understand how arterial pulsations and dilations interact with brain deformability to drive fluid flow in the PVS. In simulations with compliant brain tissue, arterial pulsations did not drive appreciable flows in the PVS. In contrast, when the artery dilated with dynamics like those seen during functional hyperemia, there was a marked movement of fluid through the PVS. Our simulations suggest that in addition to its other purposes, functional hyperemia may serve to increase fluid exchange between the PVS and the subarachnoid space, improving the clearance of metabolic waste. We measured displacement of the blood vessels and the brain tissue simultaneously in awake, head-fixed mice using two-photon microscopy. Our measurements show that brain tissue can deform in response to fluid movement in the PVS, as predicted by simulations. The results from our simulations and experiments show that the deformability of the soft brain tissue needs to be accounted for when studying fluid flow and metabolite transport in the brain.

## Introduction

The brain is surrounded by cerebrospinal fluid (CSF), and the movement of CSF can transport metabolic waste out of the brain^1–3^. The nature of CSF movement into the brain tissue (where it becomes interstitial fluid, ISF) is currently a source of controversy^4,5^. Recent work^1,6^ has suggested that CSF is actively transported along the paravascular space (PVS) around arteries. The PVS is a fluid-filled region between the arterial smooth muscle and astrocyte endfeet, and it is connected to the sub-arachnoid space (SAS). The bulk movement of CSF is thought to be driven by heart beat-driven pulsations of arteries, which push CSF into the brain (“peristaltic pumping”)^7–9^. However, several studies have given a conflicting view, suggesting that there is no bulk fluid movement, and that all transport in the brain is due to diffusion^3,10–13^.

An important approach for understanding fluid movement in the brain and the PVS is simulation of fluid dynamics. Calculations based on fluid mechanics^8,14,15^ do not agree on the magnitude and direction of the proposed “peristaltic pumping” mechanism. Moreover, the previously published models have treated the brain tissue as a rigid solid for simplicity. In reality, brain tissue is very compliant^16–19^ (‘soft’), with a shear modulus in the range of 1-8kPa. While a few models of fluid flow in the brain have considered physiological constraints on pressure differences^12^, none have sought to incorporate tissue deformation in response to the variation in pressure. Given the compliant nature of the brain^16–19^, even relatively small pressure changes will cause deformations of the tissue and would consequently produce fluid movements very different from those that would occur if the brain were rigid.

In this study, we used finite element simulations to model fluid movement in the PVS. The anatomical regions of interest in our calculations include the cerebral arteries, the PVS, the brain tissue and the SAS (Fig 1A). In our models, we have two contiguous fluid-filled compartments, namely the PVS and the SAS. Several experimental and modeling studies^10,12,13,20^ suggest that there is no appreciable fluid flow in the brain tissue and that metabolite transport in the brain occurs through diffusion. Consistent with these experimental results, we modeled the brain as a solid with no fluid flow through the tissue. To make the calculations and interpretation of results simpler, we assume a cylindrically symmetric geometry with the centerline of the artery as the axis of symmetry (Fig 1B). We performed fluid-structure interaction simulations that couple the mechanics of fluid movement in the PVS and the solid mechanics of the brain tissue. Our simulations suggest that arterial pulsations cannot drive appreciable movement of CSF in the PVS. Our simulations also suggest that neural activity-driven functional hyperemia can drive large fluid exchange between the PVS and the SAS, improving metabolite clearance from the brain. These models predicted arteriole dilation-induced deformation of the surrounding brain tissue, which we experimentally verified in awake mice. These results suggest that in addition to its involvement in other processes, functional hyperemia can drive the circulation of CSF.

**Figure 1.**
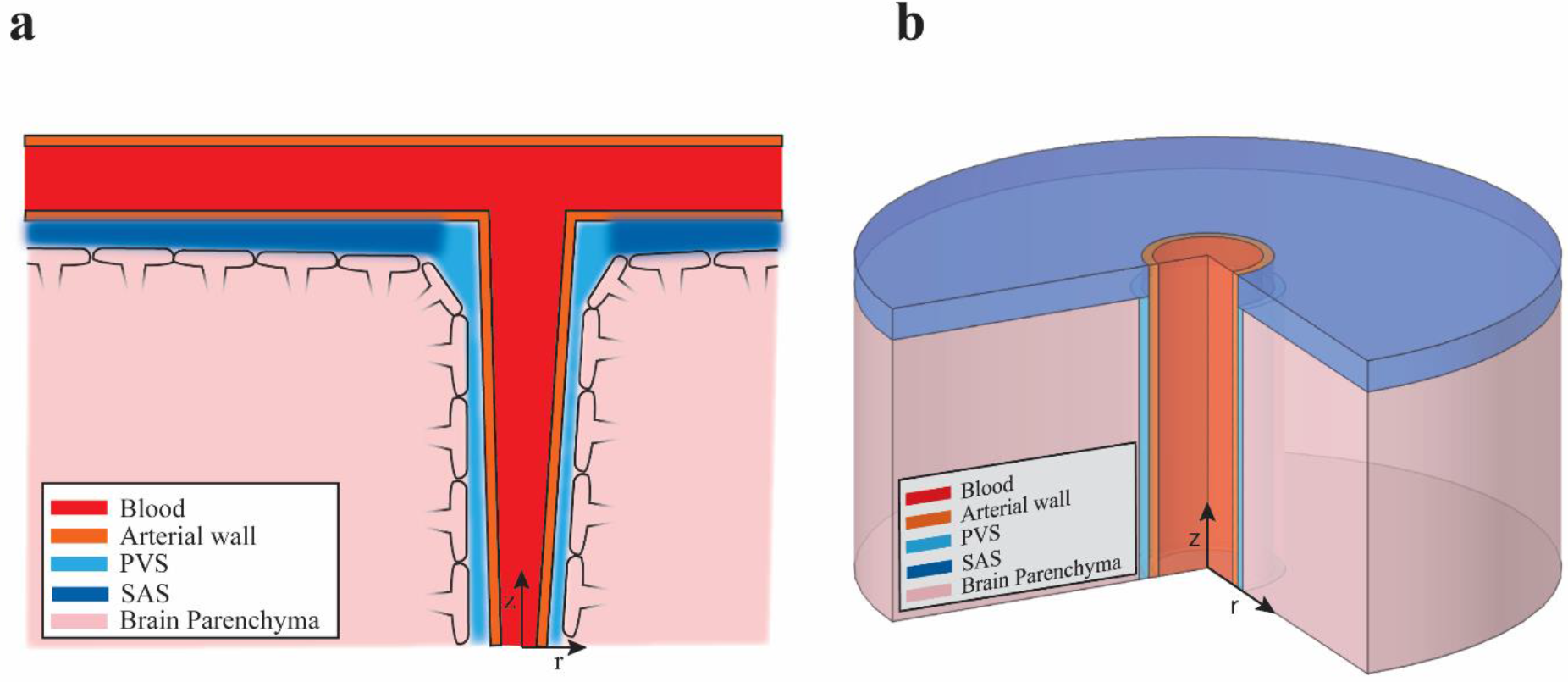
Schematic of the anatomical structure around a penetrating arteriole. **a.** Depiction the fluid filled PVS between the arterial wall and the brain parenchyma, adapted from Abbot *et al*^3^. The glia limitans covers the surface of the brain tissue and forms the brain-PVS interface. The subarachnoid space (SAS) and paravascular spaces (PVS) are interconnected fluid-filled compartments. **b.** Geometry of the computational model of a penetrating arteriole and the brain and fluid around it. The model is cylindrically symmetric around the penetrating arteriole, allowing us to use axisymmetric simulations (see appendix for full mathematical detail).

## Results

### Ignoring brain deformability leads to implausibly high pressures

We are interested in understanding how the motions of the arterial walls drive fluid flow in the PVS. We performed fluid mechanics simulations of the CSF in the PVS with the assumption that the brain is rigid. We first investigate the idea that heartbeat pulsations propagating through the arterial wall can pump CSF into the brain^8,14^ (peristaltic pumping). In this model, the space between the penetrating artery (the inner wall of the PVS) and the brain (the outer wall of the PVS) is filled with fluid. Fluid enters or exits the PVS at both the top (pial) and bottom ends of the PVS. To quantify the flow driven by peristalsis alone, we impose no pressure difference across the two ends of the PVS. We posit that fluid movement in the PVS is governed by the Darcy-Brinkman^21^ equation, which is used to simulate flow through highly porous regions^22^. Consistent with the assumption that the brain tissue is rigid, the position of the outer wall of the PVS is fixed (as was done in other models^8,15^). A detailed description of the mathematical formulation of the resulting initial-boundary value problem is provided in the appendix.

To simulate the peristaltic wave due to the heartbeat, the position of the inner wall of the PVS was prescribed via a travelling sinusoidal wave whose amplitude^9^, frequency^23^ and velocity^24,25^ were taken from experimental observations in mice. When the PVS was of anatomically realistic size (3 μm wide and 250 μm long, see methods) we observed no appreciable unidirectional pumping of fluid (0.75 μm^3^/s, 0.0015% volume of PVS per second). Instead, we see periodic fluid movement in and out of the top and bottom ends of the PVS (Fig 2b., see also Fig S1) resulting in an oscillatory flow with negligible unidirectional pumping. There is no net fluid movement in these conditions because the wavelength of the cardiac pulsation (0.1 m, see table 1) is much longer than the PVS (150-500 μm). When the wavelength of the pulsation is substantially larger than the length of the PVS, the arterial wall cannot capture the shape of the peristaltic wave. The entire length of arterial wall moves in or out almost simultaneously. This effect can be better understood by comparing the arterial wall movement in a 250 μm artery (Fig S1) with a 0.1 m artery (Fig S2).

**Figure 2.**
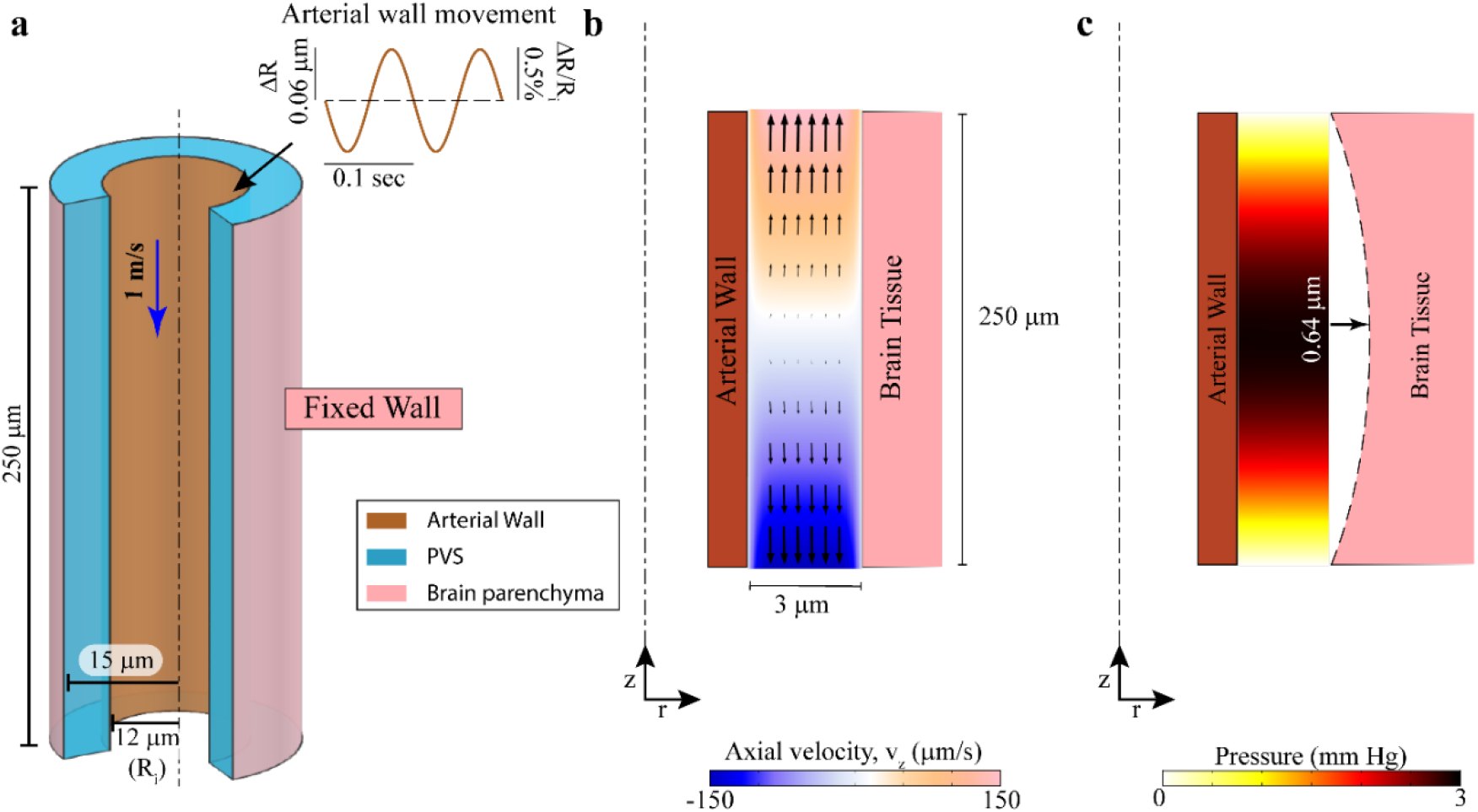
Modeling fluid flows and induced pressures in a “rigid brain” model. Note the geometry is depicted with an unequal aspect ratio in the radial (r) and axial (z) directions for viewing convenience. **a.** Geometry of PVS in in our model. The outer wall of the artery is shown in orange and the boundary of the brain parenchyma is shown in pink. For clarity, the blood and brain tissue are not shown. The dashed line represents the centerline of the artery. The inset shows the imposed heartbeat-driven pulsations in arterial radius (±0.5% of mean radius^9^,Ri) at 10 Hz, the heartrate of an un-anesthetized mouse. The pulse wave travels at 1 meter per second along the arterial wall, into the brain^25,88^ (blue arrow). In **b** and **c**, a cross section of the PVS is shown together with the surrounding arterial wall and brain tissue. **b.** Plot of the fluid velocity induced in the PVS by the arterial pulsation. Contour showing the axial velocity (velocity in the z-direction) in a cross-section of the PVS at the beginning of each sinusoidal cycle which has the highest velocity magnitudes in the pulsation cycle. (see Fig S1 for full cycle). The colors indicate the direction and magnitude of flow. Fluid velocity vectors (arrows) are provided to help the reader interpret the flow direction from the colors. The fluid flow is nearly symmetric through both ends of the PVS, so there is very little net unidirectional pumping. However, since the velocity magnitudes are high, transport of metabolites can be improved by mixing of the fluid in the PVS and the SAS^15^. Note: Arterial and brain tissue displacements induced by arterial pulsations are very small (<0.1 μm). To make the movements clearly visible, we scaled the displacements by a factor of 10 in post-processing. **c.** Fluid pressure in the PVS corresponding to the flow shown in b. Pressure changes due to fluid flow in the PVS reach several mmHg. These pressures will deform the soft brain tissue, which has a shear modulus of 1-8 kPa^86,97^ (8-60 mmHg). The dotted line shows the estimated deformation in the brain tissue (shear modulus 4kPa – Kirchhoff/De Saint-Venant elasticity with Poisson ratio of 0.45) from the pressure shown in the figure. Under these assumptions, the deformations in the brain tissue are 10 times bigger (0.64 μm) in magnitude compared the peak of heartbeat driven pulsations (0.06 μm – shown on inset in a). Therefore, the deformability of brain tissue cannot be neglected.

**Table 1.** Parameters used in simulations

| Parameter Name | Symbol | Default | Range | Unit | Source |
| --- | --- | --- | --- | --- | --- |
| Arterial radius | $R_1$ | 12 | 5 to 20 | $\mu\text{m}$ | 34,36,46 |
| PVS length | $L_a$ | 250 | 250 to 500 | $\mu\text{m}$ | 36,90 |
| PVS width | wd | 3 | 2-10 | $\mu\text{m}$ | 1,9,78 |
| CSF viscosity | $\mu_f$ | 0.001 | - | Pa.s | 79,80 |
| CSF Density | $\rho_f$ | 1000 | - | $\text{kg/m}^3$ | 79,80 |
| PVS porosity | $\zeta$ | 0.8 | 0.5-0.9 | - | 9,57,81 |
| PVS permeability | $k_s$ | $2 \times 10^{-14}$ | $7 \times 10^{-13}$ to $2 \times 10^{-15}$ | $\text{m}^2$ | 82,83 |
| Brain section radius | $R_3$ | 150 | 100-200 | $\mu\text{m}$ | 90,91 |
| Brain shear modulus | $\mu_s$ | 4 | 1-8 | kPa | 16-19,71,72,86,87 |
| Pulsation amplitude<br>(% arterial radius) | $b_1$ | 1 | 0.5-2 | - | 9 |
| Pulsation Frequency | f | 10 | 7-14 | Hz | 23 |
| Pulse wave speed | c | 1 | 0.5-10 | m/s | 25,88,89 |
| Pulse wave wavelength | $\lambda$ | 0.1 | 0.03-1.43 | m | c/f |

Our result is very similar, in terms of magnitude and direction of fluid velocities(Fig 2b), to that by Asgari *et al*^15^, who used a similar PVS geometry in their model. Asgari *et al*^15^ showed that large oscillatory fluid flow in the PVS can promote fluid mixing within the PVS and in between the PVS and the SAS and thus improve metabolite transport. When we simulated a PVS 0.1m in length, we saw pumping of fluid, consistent with Wang and Olbricht^8^, and Schley *et al*^14^ (Fig S2a). However, these models predict pressure differences of up to 2.0×10^5^ mm of Hg (Fig S2a). This is comparable to the pressures found on the ocean seabed, under several kilometers of water (2.0×10^5^ mm of Hg = 2.7 km of water), which is physically implausible.

Modeling the brain-PVS interface as fixed presumes that the brain tissue is rigid. This assumption is only valid if the pressures produced are small relative to the elastic modulus of the brain. When the brain is presumed rigid, our simulations show that the peak pressures in the PVS during pulsations can reach 3 mmHg (Fig 2c). Given that the brain is a soft tissue with a shear modulus in the range of 1-8 kPa^16–19^ (7-30 mmHg), we estimated that the peak displacement of the brain tissue induced by the pressure profile in Fig 2c would be 0.64μm (with a shear modulus of 4 kPa). This displacement value stands in stark contrast to the fact that the arterial wall displacement driving the flow is only 0.06μm. We conclude that pressures induced by the flow demand that the mechanical properties of brain tissue and its deformability must be accounted for to accurately simulate fluid dynamics.

### Arterial pulsations do not drive flow in the PVS in a compliant brain model

We modified our model by treating the brain as a compliant, elastic solid (Fig 3a). The brain tissue was modelled as a compressible, Saint-Venant-Kirchhoff solid with a Poisson’s ratio of 0.45. The pressure and the fluid shear forces in the PVS were coupled to the elastic deformation in the brain tissue using force balance equations at the interface. We coupled the fluid velocity with the velocity of deforming brain tissue, to create a fully-coupled, fluid-structure interaction model (Fig 3b). The boundary conditions in this model were different from the one used in the previous section, where both ends of the PVS were open to flow. Here, the perivascular space was capped on the brain (bottom) end, preventing fluid flow. We did this because the PVS around penetrating arteries is tapered and almost completely vanishes a few hundred micrometers into the brain, usually right before the penetrating arteriole branches into capillaries^2^. The pial opening of the PVS is connected to a flow resistance, which models fluid moving into and out of the fluid reservoir in the subarachnoid space (Fig 3a).

**Figure 3.**
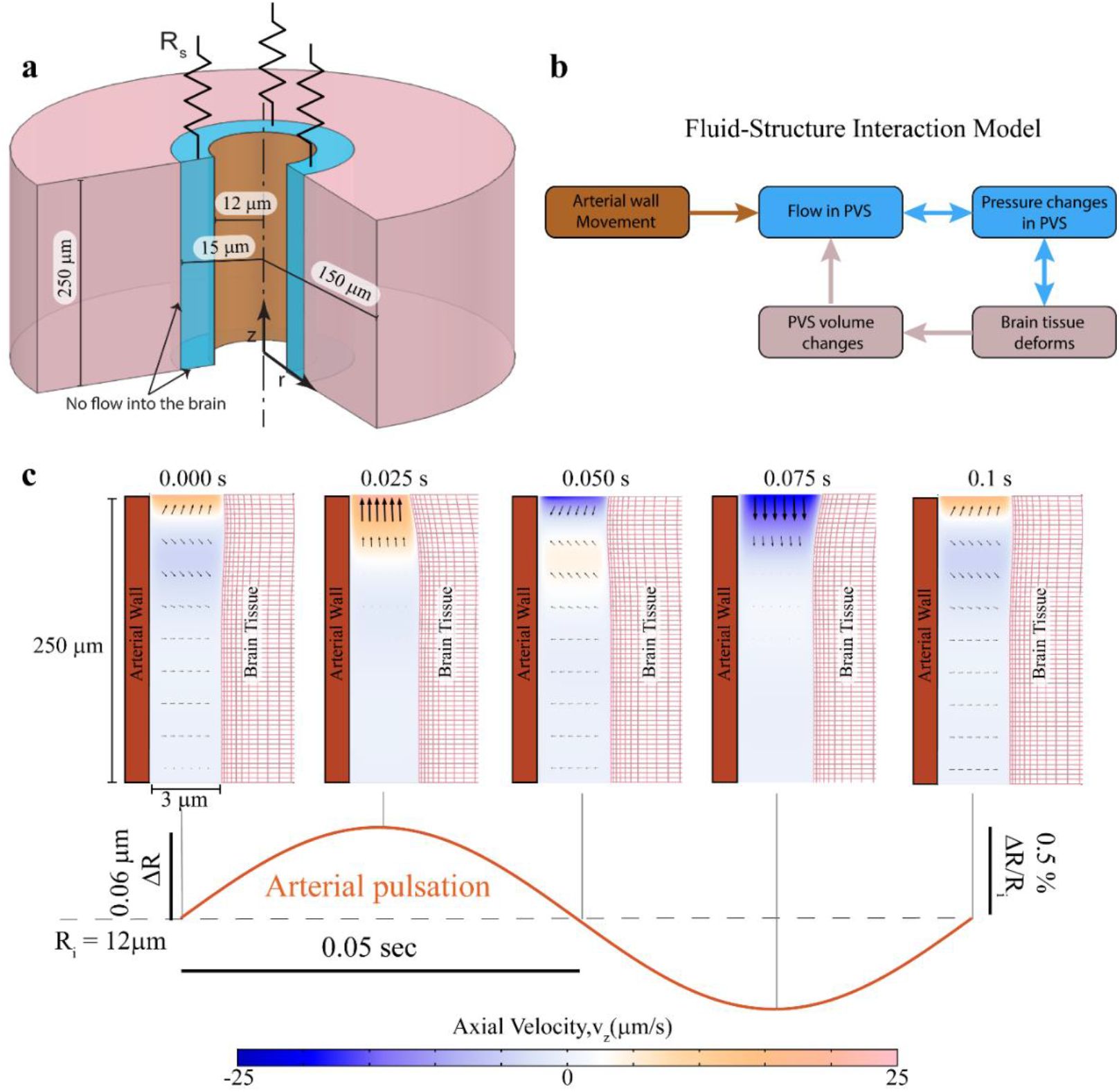
Arterial pulsations do not drive flow in the PVS in an arterial-brain model with realistic mechanical properties. Note the geometry is depicted with an unequal aspect ratio in the radial (r) and axial (z) directions for viewing convenience. a. The model of the penetrating artery. The brain tissue is modelled as a compliant solid. The subarachnoid space is modelled as a flow resistance (R_s_) at the end of the PVS. For the simulation with the subarachnoid space modelled as a fluid filled region, see Fig S4. b. A schematic depicting the fluid-structure interaction model described in a. The arterial wall movement drives the fluid movement in the PVS. This fluid movement is coupled with the pressure changes. These pressure changes deform the brain tissue, changing the shape and volume of the PVS. These volume changes will affect the flow in the PVS, as demonstrated in c. c. Plot showing the axial fluid velocity (velocity in the z-direction) in a cross section of the PVS, when the arterial wall movement is given by periodic pulsations. The amplitude and frequency of the arterial pulsations are taken to be typical values for cerebral arteries in mice. A positive velocity value (orange) means that fluid leaves the PVS and negative value (blue) means that fluid enters the PVS. Fluid velocity vectors (arrows) are provided to help the reader interpret the flow direction from the colors. The region in white has little to no flow. These plots show that there is no significant flow into the PVS driven by arterial pulsations. Note: Arterial and brain tissue displacements induced by arterial pulsations are very small (<0.1 μm). To make the movements clearly visible, we scaled the displacements by a factor of 10 in postprocessing.

We first investigated how the compliant brain tissue model would respond to arterial pulsations. We imposed movement of the arterial wall with the same dynamics used in our previous model and visualized the resulting fluid flow in the axial direction (V_z_) (Fig 3c). Throughout the pulsation cycle, most of the fluid in the PVS showed little to no movement (white). This lack of movement of fluid in the PVS in response to arterial pulsations held true over a wide range of changes in assumptions and parameters. Changing the brain tissue model from nearly incompressible (Poisson’s ratio of 0.45) to a completely incompressible(Poisson’s ratio of 0.5), Neo-Hookean model (Fig S3) had minimal impact on the pulsation-induced flow. Pulsation-driven flows were also small in simulations where the subarachnoid space (SAS) was modeled as a fluid filled, porous region connected to the PVS (Fig S4). These small flows were due to the compliance of the brain, as any pressure gradient that could generate substantial fluid movement will be dissipated on deforming the brain tissue instead.

In order to quantify the fluid exchanged between the PVS and SAS, we defined the volume exchange fraction, Q_f_, driven by arterial wall movement. The volume exchange fraction is defined as the ratio of the maximum amount of fluid leaving the PVS to the total volume of fluid in the PVS (see appendix for full mathematical description). Arterial pulsations driven by heartbeat cause a mere 0.18% (Q_f_ = 0.0018) of the fluid in the PVS to be exchanged with the SAS per cardiac cycle. These simulations show that under physiologically plausible conditions, cardiac pulsations drive a negligible amount of CSF exchange between the SAS and PVS.

### Arterial dilations during functional hyperemia can drive fluid exchange in the PVS

While cardiac pulsations are small in size, the arterial dilations that accompany increases in local neural activity are substantially larger and longer lasting. In response to increases in local neural activity, cerebral arteries can dilate by 20% or more in non-anesthetized animals^26–29^ and are the basis for the blood-oxygen-level dependent (BOLD), functional magnetic resonance imaging (fMRI)^30–33^ signal. In contrast with arterial pulsations which occur at the heart rate, these neurally-induced arterial dilation take one to three seconds to peak and last for several seconds in response to a brief increase in neural activity.

To study the flow of CSF in the PVS driven by functional hyperemia, we imposed arterial wall motion in our model that matched those observed in awake mice during a typical functional hyperemic event^34–36^ (Fig 4a). The mathematical formulation of this problem is identical to the previous simulation, with the exception that the arterial wall movement was given by a typical vasodilation profile instead of a heartbeat-driven peristaltic wave (Fig 4a). Compared to the flow driven by arterial pulsations, functional hyperemia-driven flow in the PVS had substantially higher flow velocities (Fig 4a). The fluid movement in these simulations was substantial and indicated that arterial dilations due to a single brief hyperemic event could exchange nearly half (Q_f_ = 0.4804) of the fluid in the PVS with the SAS. The simulations also suggest that the pressure changes in the PVS due to this flow will deform the brain tissue by up to 1.2 μm for an arterial dilation of 1.8 μm (Fig S6). To check the robustness of these results, we repeated this simulation with a wide range of parameters, as well as with an incompressible elastic model (Fig S7). We also modeled the SAS as a fluid filled, porous region connected to the PVS (Fig S8). In all cases, functional hyperemia-like dilations drove substantial fluid movement in the PVS. Compared to arterial pulsations, the vasodilation driven fluid exchange between PVS and SAS was two orders of magnitude higher under a wide range of model parameters (Fig S9).

**Figure 4.**
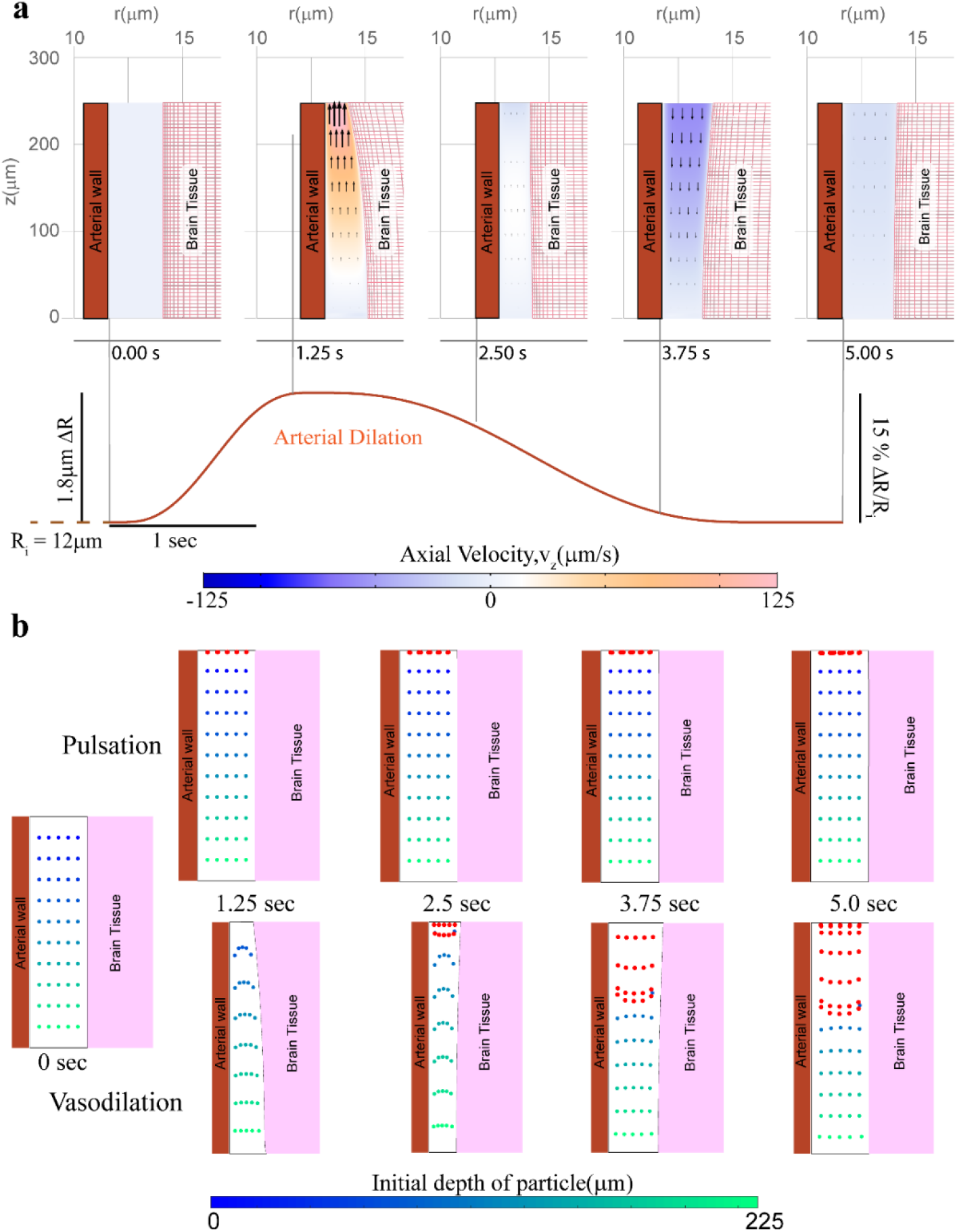
Arterial dilations during functional hyperemia can drive fluid exchange in the PVS. Note the geometry is depicted with an unequal aspect ratio in the radial (r) and axial (z) directions for viewing convenience. a. Contours showing the axial velocity (velocity in the z-direction) in a cross section of the PVS, when the arterial wall movement is given by a typical neural activity-driven vasodilation response. A physically realistic, fluid-structure interaction model (See Fig 3a) is used. Compared to heartbeat-driven pulsations (Fig 3c), vasodilation-driven fluid flow occurs through the entire length of the PVS and has substantially higher flow velocities. The model also predicts that the vasodilation can also cause significant deformation in the brain tissue. b. Comparison of particle motion in the fluid of the PVS during arterial pulsations and vasodilation. The blue-green dots represent fluid in the PVS, with the colormap showing the initial position(depth) of the fluid particle in the PVS. Fluid particles near the SAS (red dots) are added once every 0.5 secs to the simulation to simulate fluid mixing between the PVS and the SAS. There is very little fluid movement driven by arterial pulsations. Vasodilation drives appreciable fluid exchange between the PVS and the SAS.

Because the fluid movement from arterial pulsations and functional hyperemia occur at different time scales (10 Hz and 0.2 Hz, respectively), we directly compared the fluid movement driven by arterial pulsations and functional hyperemia over equal time periods. This was achieved by calculating fluid particle trajectories in the deforming geometry of the PVS (see appendix for full mathematical description of boundary value problem for particle tracking in a deforming domain). The results of these calculations indicate that a single hyperemic event can cause substantially more fluid movement in the PVS compared to arterial pulsations over the same time (Fig 4b, also see videos SV1 and SV2). The blue-green dots in Fig 4b represent fluid in the PVS, with the colormap showing the initial position(depth) of the fluid particle in the PVS. Fluid particles near the SAS (red dots) are added once every 0.5 secs to the calculation to simulate the possibility of fluid exchange between the PVS and the SAS. These calculations suggest that when the flow in the PVS is modeled with coupled soft brain tissue mechanics, functional hyperemia can drive appreciable fluid exchange between the PVS and the SAS, while arterial pulsations do not drive flow. We also simulated the flow driven by the two mechanisms over a period of 50s and found similar results (videos SV3 and SV4).

There are two main reasons why functional hyperemia drives large fluid exchange between the PVS and the SAS, while arterial pulsations are ineffective at driving fluid movement in the PVS. Firstly, heartbeat-driven changes in arterial diameter are very small (0.5-4%^9^) in magnitude compared to neural activity-driven vasodilation (10-40%^34^) and therefore there is a large difference in the volume of fluid displaced by the two mechanisms. Our measurements *in-vivo* also confirmed that the diameter changes driven by heartbeat (Fig S10) are in the 0.5-4% range while the diameter changes driven by vasodilation are in the 10-40% range (Fig 5m). A difference in the magnitude of blood volume change driven by heartbeat and hyperemia has also been observed in macaques^37^ and humans^38^ using functional magnetic resonance imaging (fMRI). Secondly, there is a large difference in the frequency of pulsations (7-14 Hz^23^ in mice, nominally 1 Hz in humans) and hyperemic (0.1-0.3 Hz^35,39^) motions of arterial walls. Fast (high frequency) movement of arterial walls cause larger changes in pressure which will deform the brain tissue and cancel the flow. Also, deformable (elastic) elements absorb more energy at higher frequencies. If the electrical circuit equivalent of flow through the PVS with a rigid brain is analogous to a resistor, the equivalent of flow through the PVS with a deformable brain is analogous to a resistor and inductor in series (Fig S11a-b). In other words, arterial wall motion at higher frequencies drives less fluid movement compared to arterial wall movement at lower frequencies. A similar phenomenon has been studied extensively in the context of blood flow through deformable arteries and veins^40–43^. We compared the fluid exchange percentage for an arterial wall movement given by a sine wave (4% peak to peak) of different frequencies, and found that the fluid exchange percentage has an inverse power law relation to frequency (f) (*Q_f_* = 2.33*f*^−0.57^ % for the default parameters, Fig S11c).

**Figure 5.**
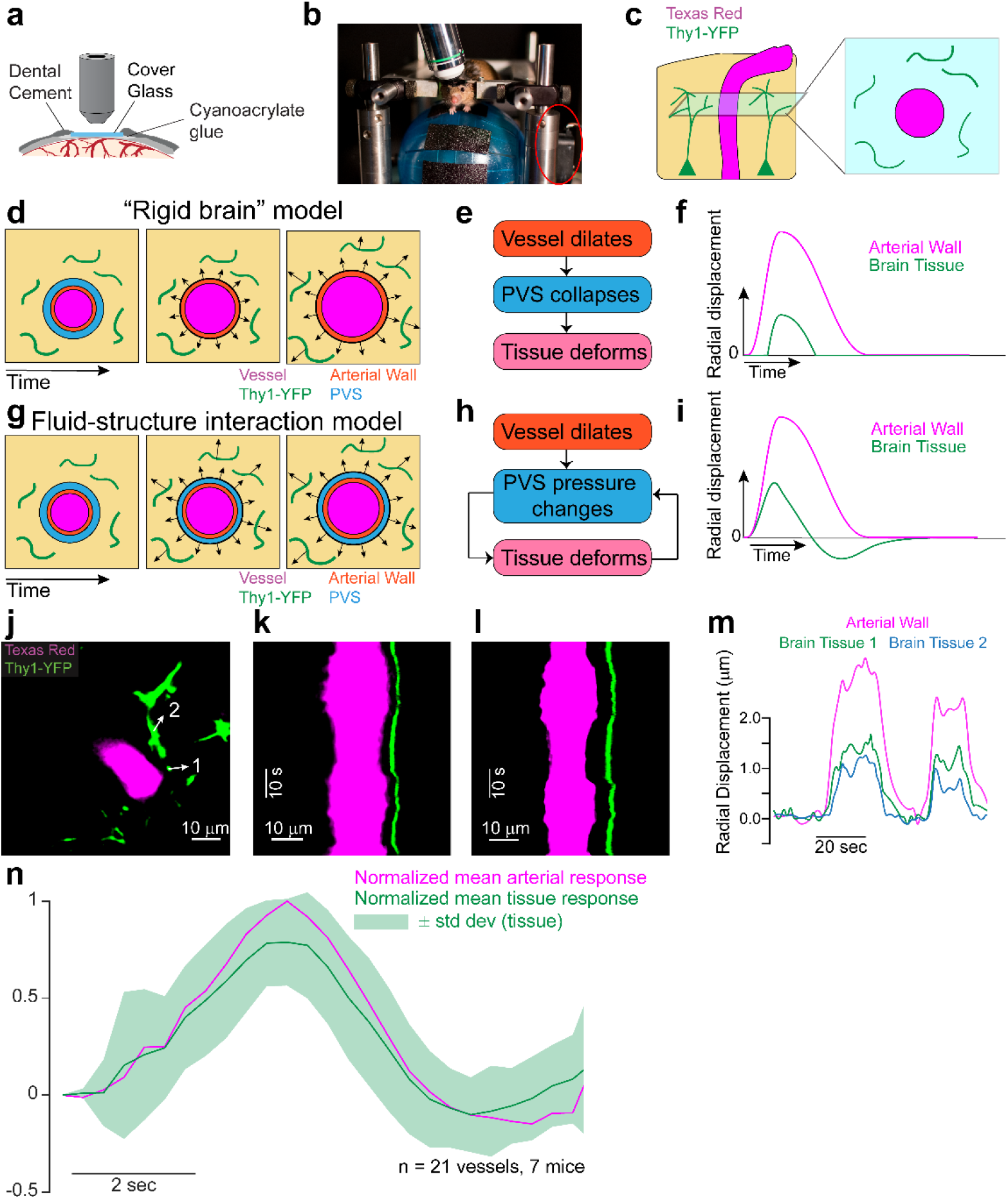
In-vivo measurement of brain tissue-displacement suggests that the brain tissue can deform because of pressure changes in the PVS. a. Schematic of a thin skulled window. Mice implanted with a thinned-skull window (PoRTS window^46^) were imaged under a two-photon laser scanning microscope(2plsm). The mice were head-fixed and allowed to run voluntarily on a spherical treadmill. b. Experimental setup for two-photon microscopy. Mice were head-fixed and placed on a spherical treadmill. The locomotion data is collected by a rotary encoder (encircled in red). c. Schematic of the fluorescent elements in the brain parenchyma (left) surrounding a penetrating artery and the expected 2-D images under a 2plsm (right). A retro-orbital injection of Texas red dye conjugated dextran (40 kDa, 2.5% w/v) makes the vessel lumen fluorescent. The yellow fluorescent protein is expressed by a sparse subset of neuronal processes. d. A schematic of the brain tissue deformations expected from a “rigid brain” model, where pressure changes in the PVS do not deform the brain. The position of the vessel wall and the PVS are shown on the left. When the artery dilates, the brain tissue would not deform until the PVS completely collapses (middle). After the PVS has collapsed completely, the brain tissue would start deforming(right). e. Flow chart of the mechanism of brain tissue deformation in a “rigid-brain” model. f. The expected radial displacement in the brain tissue in response to arterial dilation in the “rigid brain” model. The brain tissue does not deform until the PVS has completely collapsed. g. A schematic of the expected brain tissue deformation from a fluid-structure interaction model. Here the pressure changes in the PVS cause the brain tissue to deform. h. Flow chart of the mechanism of brain tissue deformation in a fluid-structure interaction model. i. The expected radial displacement in the brain tissue in response to arterial dilation in the fluid-structure interaction model (also see Fig S6). j. Median frame of the 2D image collected during in-vivo imaging. Example image of penetrating arteriole(magenta) and YFP expressing neurons(green). The arrows show the direction of the displacement measured at the location indicated by the tail of the arrow. k, l. Projection in time along a line running through the arrows 1 and 2 respectively shown in j. The images show that when the vessel dilates (indicated by a widening of the vessel in magenta), there is a corresponding radially-outward deformation in the brain tissue (indicated by the movement of the green line). Time moves forward in the in the vertically downward direction in both images. m. The calculated radial displacement in the brain tissue in response to changes in arterial radius. The data suggests that the brain tissue deforms due to pressure changes in the PVS before the PVS completely collapses. n. The average (7 mice, 21 vessels) peak-normalized impulse response of the radial displacement of the arterial wall (magenta) compared to the average peak-normalized impulse response of the radial displacement in the brain tissue (only one data point per vessel was used for this calculation). The data shows that there is no delay between displacement of arterial wall and the tissue, suggesting that the brain tissue deforms due to pressure changes in the PVS as predicted by the fluid-structure interaction model.

### In-vivo brain tissue deformation is consistent with a fluid-structure interaction model

One of the main predictions of the fluid-structure interaction model is the deformation of the soft brain tissue in response to the pressure changes in the PVS driven by arterial dilation. To test this prediction, we measured displacement of the cortical brain tissue surrounding penetrating arteries in awake, head-fixed B6.Cg-Tg(Thy1-YFP)16Jrs/J (Jackson Laboratory) mice^44^ using two-photon laser scanning microscopy^36^. These transgenic mice express the fluorescent protein YFP in a sparse subset of pyramidal neurons whose axons and dendrites are strongly fluorescent^45^. Mice were implanted with polished, reinforced thinned-skull windows^46^(Fig 5a) to avoid inflammation^47^, disruption of mechanical properties^48^ and the hemodynamic and metabolic effects^49^ associated with craniotomies. We simultaneously imaged processes of Thy1-expressing neurons and blood vessel diameters via intravenous injection of Texas-red dextran (Fig 5b). Arterioles in the somatosensory cortex dilate during spontaneous locomotion events due to increases in local neural activity^35^, so we imaged these vessels that will be naturally subject to large vasodilation. We performed piecewise, iterative motion correction of the collected images relative to the center of the artery (see Methods) in order to robustly measure the displacement of brain tissue during arterial dilations. We verified the measured brain tissue displacements

We considered two possible paradigms of brain deformation, a “rigid-brain” model and a fluid-structure interaction model. We predict the two paradigms to yield completely different results in terms of the displacement of the brain tissue observed in-vivo. In the rigid-brain model, the brain tissue will be unaffected by pressure changes in the PVS. In this model, pulsations and small dilations of brain tissue would cause flow in the PVS but no displacement of the brain tissue (Fig 5c). Only after the arterial wall comes in contact with the brain tissue (and the PVS has fully collapsed), arterial dilation would cause tissue displacement (Fig 5d). Therefore, displacement in the brain tissue in this model would be either non-existent (for small dilations), or similar to a “trimmed” version of the displacement of the arterial wall (Fig 5e). Alternatively, in the fluid-structure interaction model, any movement of the arterial wall that can drive fluid flow in the PVS will result in pressure changes in the PVS that are sufficient to deform the ‘soft’ brain tissue, as predicted by our simulations (Fig 5f, 5g). Therefore, displacement should be observed in the brain tissue as soon as the arterial wall starts to dilate. In the fluid-structure interaction model, the radial displacement in the brain tissue would be a scaled version of the radial displacement of the arterial wall (Fig 5h).

We calculated the radial displacement of the arterial wall and the brain tissue in-vivo (n = 21 vessels, 7 mice) using two-photon microscopy. The radial displacement of the brain tissue was between 20-80% of the radial displacement of the arterial wall. The simulations suggest that such a variation is to be expected due to possible changes in the width and depth of the PVS and the distance of the plane of imaging from the surface of the brain (Fig S6a and S6b). Despite the variation in the amplitude of displacement in the tissue, our simulations predict that the waveform of the displacement in the tissue should be very consistent. In particular the peak-normalized displacement response of the brain tissue should almost identical everywhere (Fig S6c). We used this result from the simulation to test the predictions of the model experimentally. We calculated the peak normalized impulse response of the displacements to locomotion (Fig 5n). The calculations of tissue displacement for each artery (an example is shown in Fig 5j-m), as well as the normalized impulse response for the brain tissue (Fig 5n) suggest that the displacement in the brain tissue started as soon as the arterial dilations started. This implies that the brain tissue can deform due to pressure changes in the PVS, as predicted by the fluid-structure interaction model. All the displacement values in the brain tissue used for calculating the average waveform reported in Fig 5n were subject to a rigorous set of tests (see methods) to account for motion artifacts. To visualize the brain tissue displacements accompanying vasodilation, we plotted a kymogram taken along diameter line bisecting the arteriole and crossing neural processes (Fig 5k, 5l). Distance from the center of the arteriole is on the x-axis and time on the y-axis. Dilations appear as a widening of the vessel, while displacements will show up as shifts on the x axis. This visualization was used as an additional step in validating the displacement values calculated by our method. For calculating the average waveform of tissue displacement shown in Fig 5n, only one of the calculated displacement values per vessel that could also be visually verified was used. The displacement of the brain tissue is also apparent from visualizing the data. Supplementary video SV5 shows 10 seconds of imaging data, where we can observe the brain tissue(green) deforms in response to dilation of the vessel(magenta).

The fluid-structure interaction model incorrectly predicted a negative radial displacement in the brain tissue, when the artery constricts or returns to its original process. This anomaly can be explained by the fact that the fluid-structure interaction model neglects the elastic forces in the connective tissue (extracellular matrix) in the PVS. Connective tissue can have a highly non-linear elastic response when the loading is changed from compression to tension. The elastic modulus of connective tissue under tension can be 2-3 orders of magnitude higher than the elastic modulus in compression^50–52^. Connective tissue is made up of networks of fibers and the energy cost of bending these fibers is several orders of magnitude smaller than stretching them. When the artery dilates, these fibers are subject to a compressive loading and they buckle(bend) rather than compress, and as a result generate very little elastic forces (Fig S11b). On the other hand, when the artery constricts or returns to its initial size, these fibers are subjected to a tensile load (Fig S11c) and produce significantly higher (2-3 orders of magnitude higher) elastic forces. However, our model only considers the fluid-dynamic forces in the PVS and neglects the elastic forces. This is one of the shortcomings of our model, that can be corrected in the future using models of poroelasticity^53–55^.

## Discussion

While there have been several models of the fluid mechanics in the PVS^8,14,15^, none of them considered the impact of the soft, deformable brain tissue on CSF flow in the PVS. Our simulations show that fluid flow in a porous PVS cannot be studied without considering of the deformability of the brain tissue. We have also presented empirical evidence to support our claim that the brain tissue deforms in response to pressure changes in the PVS. As far as we know, this is the first study to include deformability of the brain tissue in modeling fluid flow in the PVS and to use experiments to support the predictions of fluid dynamic simulations.

We have made some simplifying assumptions in our model and we see room for further improvement of the model. Firstly, we assumed a cylindrically symmetric geometry for our model. In reality, the PVS around penetrating vessels can be eccentric and elliptical^9^. An eccentric annular region with a pulsating inner wall can cause a slow drift in the fluid particles^56^ (However, the drift caused by eccentricity is not unidirectional and the bulk movement of the particles is much smaller compared to the oscillations^9,57^). Secondly, we neglected the possibility of fluid flow within the brain parenchyma and the effect of the aquaporin-4 channels found on the astrocyte endfeet lining the brain-PVS interface^3^. This can be rectified in future studies by using models of poroelasticity^53–55^, which simultaneously simulate fluid movement through the extracellular space and the deformations in the brain tissue.

Our results are in agreement with the findings of several experimental studies^9,10,20,57–59^, though they cast the results in a new light. Some studies have used microspheres to visualize the CSF movement in the PVS of pial arteries^9,57^. Similar to their results, our simulations suggest that CSF oscillates with the frequency of heartbeat driven pulsations near the surface of the brain (Fig 3c, Figures S3 and S4). It can also be shown that larger arterial pulsations can cause larger oscillations in CSF flow, similar to the case of induced hypertension found by Mestre *et al*^9^. In contrast to the conclusions of these particle tracking studies^9,57^, our simulations suggest that arterial pulsations do not provide a driving force for unidirectional pumping of CSF. This lack of unidirectional pumping is consistent with molecular weight-dependent transport of dyes in CSF observed with intra-parenchymal tracer injection in mice^10^ and intrathecal injections in rats^20,60^. Our results also agree with the findings that voluntary running, which increases neural activity^61,62^ and induces functional hyperemia^39,63^ in several regions of the brain, enhances penetration of tracers injected into the cisterna magna^58^. The silencing of neural activity (and therefore vascular activity) by anesthetics^64,65^ can explain diminished movement of tracers injected into the cisterna magna under anesthesia^66^. The variability in results between groups may be influenced by anesthesia type and levels, both of which have large effects on the amplitude of the arterial dilations elicited during functional hyperemia^34^. Finally, brain-wide hyperemia observed during sleep^67^ can explain improved tracer transport in the brain observed during sleep^59^.

Our results have implications for the development and treatment of CNS disorders and suggest that in addition to its other purposes, functional hyperemia may serve to improve transport in the brain. Several studies support the idea that vascular dysfunction can be a precursor to neurodegenerative diseases^68–70^. Our simulations suggest a mechanistic relation between neurovascular coupling and metabolite clearance from the brain, which can explain the development of neurodegenerative diseases like Alzheimer’s. The response of our model to changes in key parameters can explain the effect of aging on clearance of metabolic waste from the brain. Some studies have shown that the elastic modulus of the brain decreases with aging^71,72^, and our model predicts less fluid exchange between the SAS and the PVS when the elastic modulus is lowered (Fig S9a). Finally, increase of PVS width with aging^73^ might be a reason for reduced clearance of metabolic waste from the brain (Fig S9b).

## Methods

### Modeling assumptions

The simulations were performed with the assumption of axisymmetric geometry (Fig 1). The flow in the PVS was modeled as Darcy-Brinkman flow through a highly porous region. Arterial wall displacements, and later brain tissue deformability, cause the PVS to be a time-dependent domain. To properly account for the motion of the PVS we adopted an arbitrary Lagrangian-Eulerian approach^74^ (ALE). The motion of the PVS (often referred to as a mesh motion or ALE map) was modeled using a harmonic model^75^. The ALE implementation ensures that the fluid dynamics are not affected by the choice of the model for the deformation of the fluid-filled region. The brain tissue was modeled as an elastic solid. Unless otherwise stated, we modeled the brain as a compressible, De Saint-Venant-Kirchhoff material with a Poisson’s ratio of 0.45. For most of the simulations, the pial opening of the PVS was connected to a flow resistance, which models fluid moving into and out of the subarachnoid space. The flow resistance was implemented as a Robin boundary condition, i.e., a flowrate-dependent pressure-like traction was applied at the pial opening of the PVS. Simulations where the subarachnoid space (SAS) was modeled as a fluid filled, porous region connected to the PVS (Fig S4, S8) confirmed that the Robin boundary condition^76^ is adequate to simulate the flow resistance of the SAS.

The interaction between the flow in the PVS and the elastic deformation of the brain was implemented using a fluid-structure interaction model. The displacement of the PVS at the Brain-PVS interface is made equal to the displacement of the brain tissue. Similarly, the velocity of the fluid at the Brain-PVS interface is made equal to the velocity of the brain tissue. The forces from the fluid flow (pressure and fluid shear) in the PVS are applied as a boundary force on the brain tissue at the Brain-PVS interface. This coupling of displacements, velocity and forces is implemented simultaneously to create a fully coupled fluid-structure interaction model^77^.

### Model Parameters

We chose parameters for the model based on previous published experiments in adult mice (see table 1). Arterial radius was varied between 5-20 μm^34,36,46^. Because the PVS becomes very small in the range of 150-500 μm from the pial surface^3^, the PVS was modelled as a straight section a length of 150-500 μm. The width of the PVS was between 2-10 μm^1,9,78^. The viscosity (0.001 Pa*s) and density (1000 kg/m^3^) of CSF were taken from experimentally determined values^79,80^.

The PVS surrounding penetrating arteries is a fluid filled region with a higher porosity than the brain tissue^1,3^. However, the volume of the PVS is not entirely occupied by fluid, as evidenced by the presence of fibroblast-like cells^81^ in the PVS and impermeability to large (1 μm diameter) particles^9,57^. Therefore, the porosity (fraction of fluid volume to the total volume) of the PVS was assumed to be between 0.5-0.9. The fluid permeability of the PVS is taken from a range of possible values. The maximum value of the permeability was 7×10^−13^ m^2^, a value at which the flow resistance of the PVS is twice as much as a channel of the same dimensions with infinite permeability (see Fig S13). The minimum possible value of PVS permeability was 2×10^−15^ m^2^, the measured permeability of the brain tissue^82,83^. The flow resistance of the SAS was taken to be 1% of the flow resistance of the PVS. In the models where the SAS is simulated as a fluid filled region connected to the PVS, the fluid permeability of the SAS was assumed to be a factor of 10 greater that of the PVS.

The radius of the simulated section of brain tissue was taken to be in the range of 100-200 μm, half of the typical distance between two penetrating arteries in the mouse cortex^84,85^. The elastic (shear) modulus of the brain tissue is taken to be between 1-8 kPa, spanning the values found in the literature^16–19,71,72,86,87^.

Heartbeat drives changes of 1-4% (peak to peak) in the radius of pial arteries in mice^9^. These pulsations travel at a speed of 0.5-10 m/s along the arterial tree^25,88,89^. Mice have a heartrate of 7-14 Hz when they are awake and freely behaving^23^. Neural activity can drive 10-30% changes in arterial radius. These changes take place at a nominal frequency range of 0.1–0.3 Hz.

### Model implementation

All the partial differential equations that govern the physics of the problem were implemented using the Galerkin finite element method^76^. All the finite element simulations were performed using COMSOL Multiphysics^98^. We used the Weak Form PDE interface (pde stands for partial differential equation) in the Mathematics Module in COMSOL to implement the governing equations on an axisymmetric geometry. The strong form of the vector equations for each problem are given in the appendix. The equations are converted to a weak form in an axisymmetric (r,z) coordinate system using Wolfram Mathematica^100^. A backward difference formula (BDF) scheme was used for the time-dependent problems in COMSOL.

The particle trajectories in Fig 4b and supplementary videos are estimated by calculating the fluid velocities, as observed from the mesh coordinates (which themselves change with time) in COMSOL. We then export these velocity values along with the corresponding mesh displacement values. These values are taken into MATLAB^99^, where we implemented a script to calculate fluid particle trajectories using the forward-Euler time integration scheme^76^.

### Surgical procedures

All procedures were performed in accordance with protocols approved by the Institutional Animal Care and Use Committee (IACUC) of Pennsylvania State University. Mice were anesthetized with isoflurane (5% induction, 2% maintenance) for all surgical procedures. The scalp was resected, and the connective tissue removed from the surface of the skull. A custom-machined titanium headbar (https://github.com/KL-Turner/Mouse-Head-Fixation) was affixed with cyanoacrylate glue (32402, Vibra-Tite) immediately posterior to the lambda cranial suture. Three self-tapping, 3/32” #000 screws (J.I. Morris) were implanted into the skull, one in each frontal bone, and one in the contralateral parietal bone. A ~4mm x ~5 mm polished and reinforced thinned-skull window was implanted over the right hemisphere somatosensory cortex as previously described^36,46^. After thinning, the skull was polished with 4F and 3F grit, and a #0 glass coverslip (Electrode Microscopy Sciences, #72198) was attached to the thinned portion of the skull with cyanoacrylate glue. Dental cement (Ortho-Jet) was used to seal the edges of the window and connect the headbar and headscrews. At the conclusion of the surgery the mice were returned to their home cage and allowed 2 days of recovery before being habituated to head fixation. Mice were habituated to head-fixation on a spherical treadmill (60 mm diameter) for 2-3 days before imaging. The mice were head-fixed for 30 mins on the first session and the length of the session was increased to 90 mins on the final session. The mice were monitored for any signs of distress during the period of habituation.

### Two-photon imaging

Prior to imaging, the mice (n=7) were briefly anesthetized with isoflurane and retro-orbitally injected with 50μL of 2.5% w/v of Texas-red conjugated dextran (40 kDa; Sigma-Aldrich), then head-fixed upon a spherical treadmill. The treadmill was coated with a slip-resistant tape and connected to a rotary encoder (US Digital, E7PD-720-118) to monitor changes in velocity of the treadmill. The changes in velocity (acceleration) were used to identify periods of rest and motion. Images were collected under a Sutter moveable objective microscope with either a 16x 0.8 NA objective or a 20×1.0 NA objective (Nikon). A MaiTai HP laser tuned to 920 nm was used to excite the YFP and the Texas-Red. The power exiting the objective was between 30-70 mW. Arteries were visually identified by their more rapid blood flow, rapid temporal dynamics of their response to locomotion, and vasomotion^23,34,92^. A two-channel photomultiplier setup was used to collect fluorescence from YFP and Texas-red. Images were collected at a nominal frame rate of 3-8 Hz.

### Data processing

A detailed flow chart of the procedure used for data processing is given Fig S14. All the data analysis was performed using MATLAB^99^ except for the visual verification of displacement, which was performed using ImageJ (NIH). All the code is available on GitHub (https://github.com/kraviteia89/Thy1-displacement).

We used the red channel for motion correction (registration using discrete Fourier transform^93^) to remove movement in order to generate movies where the center of the vessel was fixed. A 3D median filter (3,3 pixels in space and 5 frames in time) was used to remove shot noise. Due to crosstalk, the images on the red channel contained some YFP fluorescence (brain tissue) in addition to the Texas Red (vessel lumen) signal. To remove this crosstalk, we used linear model to remove the YFP signal from the red channel.

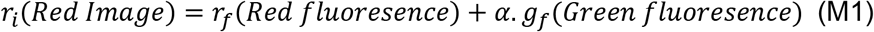

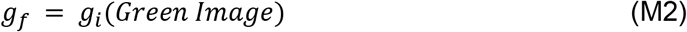

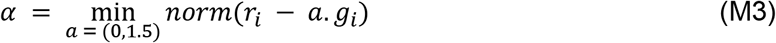

Here, the image in the red channel, r_i_, was assumed to be a linear combination of the actual red fluorescence, r_f_, and a fraction, α, of the green fluorescence, g_f_ (equation M1). The green image, g_i_, was assumed to be the actual representation of the green fluorescence (equation M2). A linear constant (α) was found, that minimized the total error under an inverse model, using MATLAB’s *fminsearch* function (equation M3). We then used equation M1 to calculate the red fluorescence r_f_.

We then estimated the changes in arterial diameter using the image sequence in the red channel. The section of the image containing the artery was cleaned-up using thresholding in radon space^94^. A rectangular region containing the artery was manually selected. The region was transformed into radon space for angles between 0° and 180° at 1° increments. At each angle, the radon transform value was rescaled between 0 and 1 to obtain a normalized value. A threshold value of 0.2 was chosen and every value below this was set to zero. We then calculated the inverse radon transform of the normalized values into the image space. The area of the vessel was calculated from the inverse-transformed image using the *regionprops* function in MATLAB. The velocity data (collected from the rotary encoder on the spherical treadmill) was used to create a binary vector of movement/rest during each frame^39^. The haemodynamic response function between the binarized locomotion and the vessel diameter was calculated by fitting the parameters of a gamma distribution function^95^. Only data sets where the goodness of fit (R^2^) between the measured vessel data and the HRF-convolved function was > 0.6 were used.

The displacement of brain tissue was calculated using a piecewise rigid motion model. A reference frame was chosen by averaging 10-30 seconds of data when the mice were resting (not moving). The images from the green channel were broken down into overlapping boxes of 64×64 pixels Consecutive boxes were 16 pixels apart in either x or y direction. The boxes containing no useful information were not used in calculating displacements. This was done by looking at the peak fluorescence in each box and only using boxes that were in the top 20 percentile of peak fluorescence. For each box, the displacement was calculated using image registration^93^ with the corresponding box in the reference frame. We used an iterative approach to calculate the displacements, meaning that each box was displaced by the negative value of the calculated displacement and the displacement between the reference and the corrected box was recalculated. This process was repeated for 5 iterations. The calculated displacement value was accepted only if the displacements converged, *i.e*., the displacement calculated in the last iteration was smaller than 1% of the total calculated displacement. This criterion was necessary because we observed several instances of movement of the brain tissue out of the plane of imaging. The DFT registration algorithm gives out a displacement value even when the reference and target images do not match. An example of the iterative method at work is shown in Fig S14b. Additionally, we use a threshold in the error (<70%) calculated by the DFT registration to accept or reject the calculated displacement. These above-threshold points were scrubbed and a median filter was used to fill in the scrubbed data points. A wavelet-based filter was used to denoise the resulting time series. A wavelet-based filter with a “biorthogonal 3.3” wavelet was used because it was found to be most efficient in extracting gamma-function like signal from noise.

Only datasets of the calculated displacement time-series that met certain criteria were included. Firstly, the direction of calculated the displacement should be radially outward (±30°) from the centerline of the vessel. Secondly, the displacement time-series should be well correlated with the time-series of vessel diameter changes (Pearson correlation coefficient >0.8). We found that the Pearson correlation coefficient between the vessel response and tissue response in both paradigms using pseudo data was always greater than 0.85 for signal-to-noise ratio between 2-50dB. Finally, we verified that the calculated displacement was visible in a projection of the image stack in time along the line of the calculated displacement. This last step was carried out in ImageJ (NIH). It is important to note that the displacements expected in both the “rigid-brain” model and the fluid-structure interaction model meet all three criteria.

To validate our code, we tested our method on pseudo-data generated using MATLAB. We generated a random 2D array of lines oriented in different directions. We displaced the generated image uniformly radially outward with the temporal dynamics of the radial displacement given by a gamma distribution function. Varying levels of noise were added to the data to determine the robustness of the algorithm. We found that the displacements extracted by our method agree well with the input displacement, and were robust to high levels of noise (Fig S15).

We used the displacements calculated from all the datasets (n=21 vessels, 7 mice) to estimate an average peak-normalized displacement response to running (Fig 5o). For each of the datasets, we calculated the impulse response of the radial displacement of the arterial wall to locomotion events using the method of deconvolution^96^. These impulse response functions were aligned so that the peak occurred at the same time and normalized to the peak value (L-infinity norm). We then calculated the impulse response functions for the radial displacement of brain tissue and applied the same time-lag as the corresponding arterial wall motion. We normalized the brain tissue displacements by their peak values. We plotted the average, normalized radial displacement of the arterial wall and the brain tissue to consider the possibility of a “rigid-brain” model or a fluid-structure interaction model.

## Acknowledgements

This work was supported by NSF Grant CBET 1705854.

## Supplementary Figures

**Figure S1.**
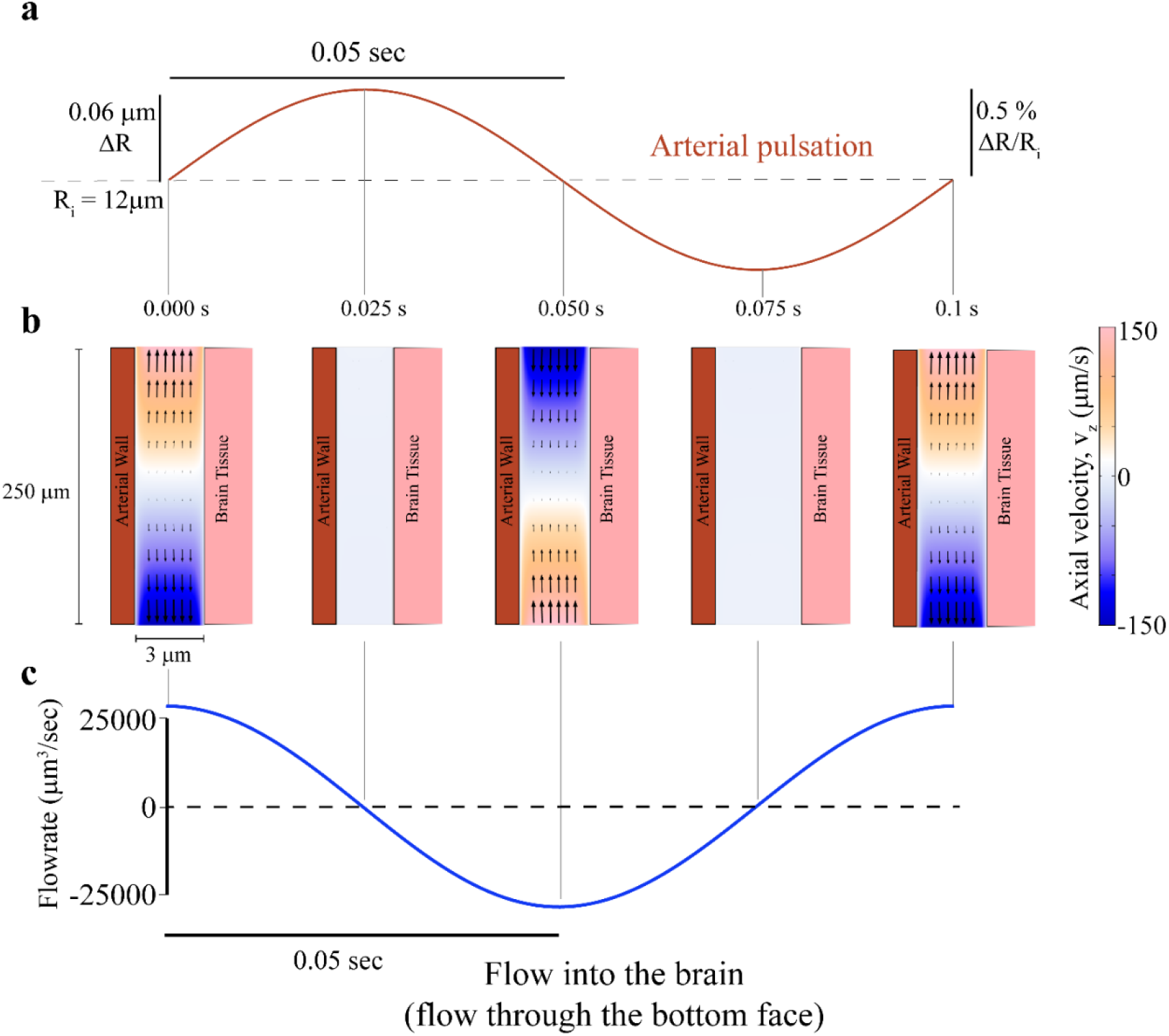
Fluid flow in a rigid brain model with realistic PVS geometry shows no appreciable unidirectional pumping. Note the geometry is depicted with an unequal aspect ratio in the radial (r) and axial (z) directions for viewing convenience. **a.** Plot of the imposed heartbeat-driven pulsations in arterial radius (±0.5% of mean radius^1^,R_i_) at 10 Hz, the heartrate of an un-anesthetized mouse. The pulse wave travels at 1 meter per second along the arterial wall, into the brain^2,3^. **b.** Plot of the fluid velocity induced in the PVS by arterial pulsation in the rigid (non-deformable) brain model. Color in the PVS shows the axial velocity (velocity in the z-direction) in a cross section of the PVS throughout the pulsation cycle. Fluid velocity vectors (arrows) are provided to help the reader interpret the flow direction from the colors. The fluid flow is nearly symmetric through both ends of the PVS when averaged over on pulsation cycle. Note: Arterial and brain tissue displacements induced by arterial pulsations are very small (<0.1 μm). To make the movements clearly visible, we scaled the displacements by 10 times in post-processing. **c.** Plot of fluid flux through the bottom surface of the PVS (into the brain) induced by the peristaltic movement of the arterial wall. The net flow over a single pulsation cycle is only 0.75 μm^3^/sec(0.0015% volume of PVS per second), while the maximum instantaneous flow rate can be as high as 25000 μm^3^/sec.

**Figure S2.**
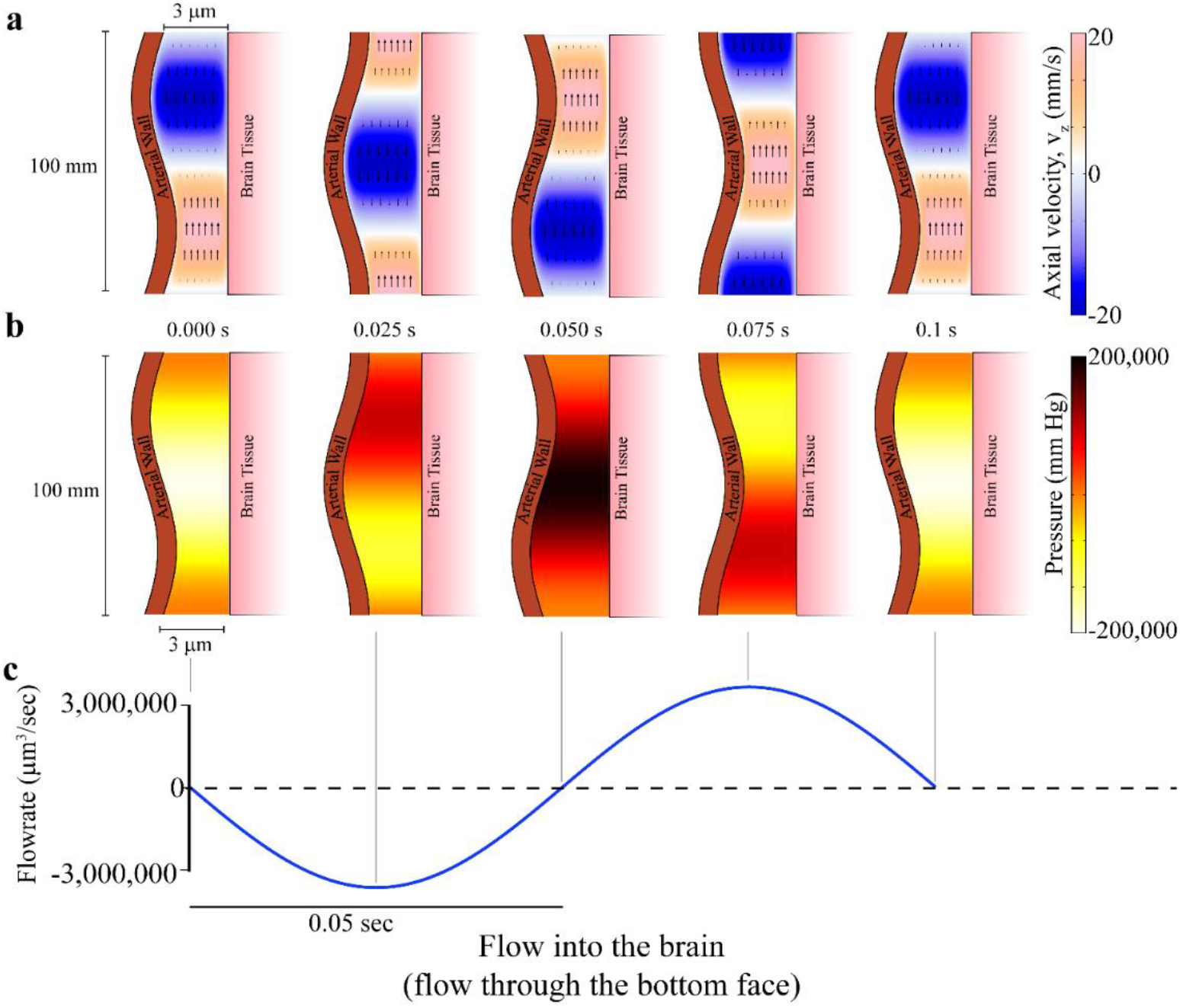
Models with unrealistic dimensions of the PVS support pumping by peristaltic movement of arteries. However, these models predict physiologically impossible pressure changes in the PVS. Note the geometry is depicted with an unequal aspect ratio in the radial (r) and axial (z) directions for viewing convenience. **a.** Plot of the fluid velocity induced in the PVS by arterial pulsation in the rigid (non-deformable) brain model, where the length of the PVS is equal to one wavelength of the peristaltic wave(0.1m, see Table 1). Color in the PVS shows the axial velocity (velocity in the z-direction) in a cross section of the PVS throughout the pulsation cycle. Fluid velocity vectors (arrows) are provided to help the reader interpret the flow direction from the colors. Note: Arterial and brain tissue displacements induced by arterial pulsations are very small (<0.1 μm). To make the movements clearly visible, we scaled the displacements by 10 times in postprocessing. **b.** Plot of the pressure induced in the PVS by arterial pulsation in the rigid (non-deformable) brain model, where the length of the PVS is equal to one wavelength of the peristaltic wave(0.1m, see Table 1). No pressure is applied at both ends of the PVS. Color in the PVS shows the pressure in a cross section of the PVS throughout the pulsation cycle. **c.** Plot of fluid flux through the bottom surface of the PVS (into the brain) induced by the peristaltic movement of the arterial wall. The net flow over a single pulsation cycle is 36,438.7 μm^3^/sec (71.6% volume of PVS per second), while the maximum instantaneous flow rate can be as high as 3,000,000 μm^3^/sec. The predictions of this model cannot be trusted because it uses an unrealistic length of the PVS and predicts impossible pressure changes.

**Figure S3.**
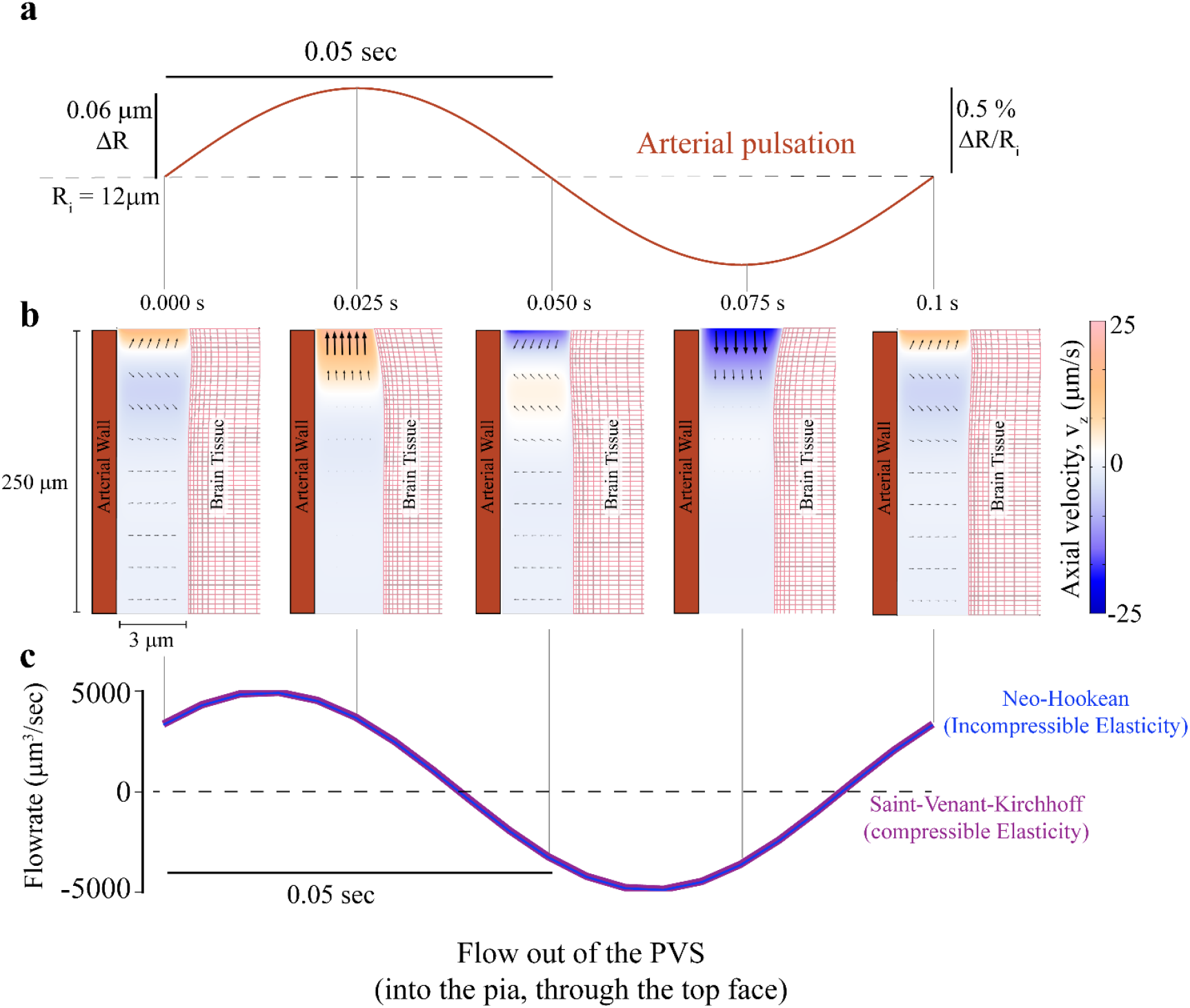
Pulsation-induced fluid flows in the PVS are small in an incompressible Neo-Hookean brain model. Note the geometry is depicted with an unequal aspect ratio in the radial (r) and axial (z) directions for viewing convenience. **a.** The imposed heartbeat-driven pulsations in arterial radius (±0.5% of mean radius^1^,R_i_) at 10 Hz, the heartrate of an un-anesthetized mouse. The pulse wave travels at 1 meter per second along the arterial wall, into the brain^2,3^ **b.** Colors showing the axial velocity (velocity in the z-direction) in a cross section of the PVS, when the arterial wall movement is given by periodic pulsations. A positive velocity value (orange) means that fluid leaves the PVS and negative value (blue) means that fluid enters the PVS. Fluid velocity vectors (arrows) are provided to help the reader interpret the flow direction from the colors. The white region is stationary. These plots (compare to those in Fig 3c) show that there is no significant flow into the PVS driven by arterial pulsations. Note: Arterial and brain tissue displacements induced by arterial pulsations are very small (<0.1 μm). To make the movements clearly visible, we scaled the displacements by 10 times in post-processing. **c.** Flow out of the PVS and into the subarachnoid space, through the pial opening of the PVS. The flow rates predicted by the model with nearly incompressible (Poisson’s ratio of 0.45) (magenta) and a completely incompressible, Neo-Hookean models (blue) were nearly identical.

**Figure S4.**
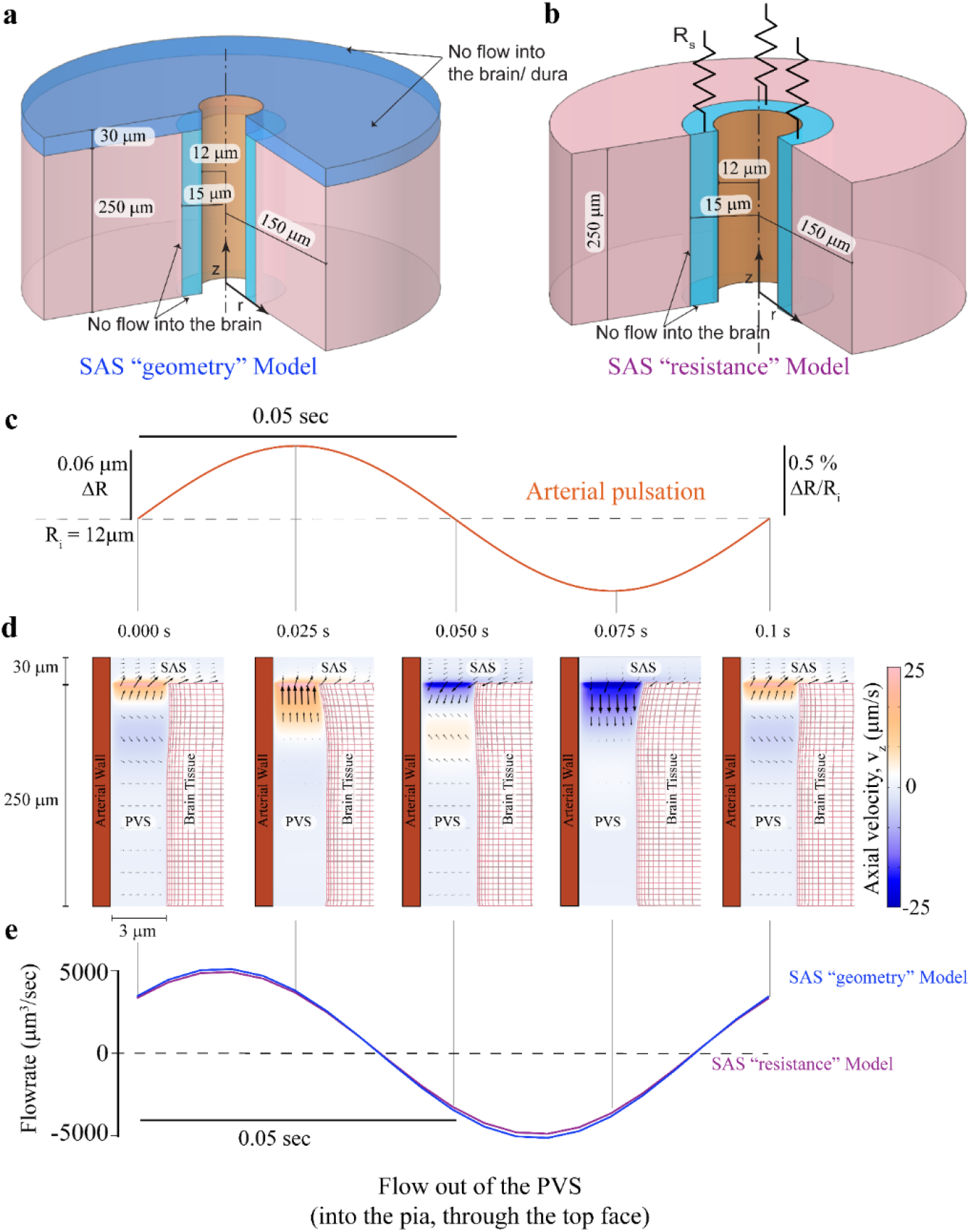
Pulsation-driven flows are small in simulations when the subarachnoid space (SAS) is modeled as a porous, fluid-filled region. Note the geometry is depicted with an unequal aspect ratio in the radial (r) and axial (z) directions for viewing convenience. **a.** Schematic showing the model of the penetrating artery used in this simulation. The brain tissue is modelled as a compliant solid. Subarachnoid space is modelled as a fluid filled region (SAS “geometry” model). **b.** Schematic showing the alternative model of the penetrating artery (Same as Fig 3a). The Subarachnoid space is modelled as a flow resistance (R_s_) at the end of the PVS (SAS “resistance” model). The results for the SAS “resistance” model are shown in Fig 3. **c.** The imposed heartbeat-driven pulsations in arterial radius (±0.5% of mean radius^1^,R_i_) at 10 Hz, the heartrate of an un-anesthetized mouse. The pulse wave travels at 1 meter per second along the arterial wall, into the brain. **d.** Plot showing the axial velocity (velocity in the z-direction) in a cross section of the PVS and the connected SAS, when the arterial wall movement is given by periodic pulsations. A positive velocity value (orange) means that fluid leaves the PVS and negative value (blue) means that fluid enters the PVS. Fluid velocity vectors (arrows) are provided to help the reader interpret the flow direction from the colors. Because the fluid is incompressible, the flow speed decreases when flowing into the SAS, which has a larger area of cross section compared to the PVS. The region in white has little to no flow. These plots show that there is no significant flow into the PVS driven by arterial pulsations. Note: Arterial and brain tissue displacements induced by arterial pulsations are very small (<0.1 μm). To make the movements clearly visible, we scaled the displacements by 10 times in post-processing. **e.** Plot of the fluid flow through the top face of the PVS into the SAS. The flow rates predicted by the SAS “resistance” model (magenta) and the SAS “geometry” model (blue) are nearly identical.

**Figure S5.**
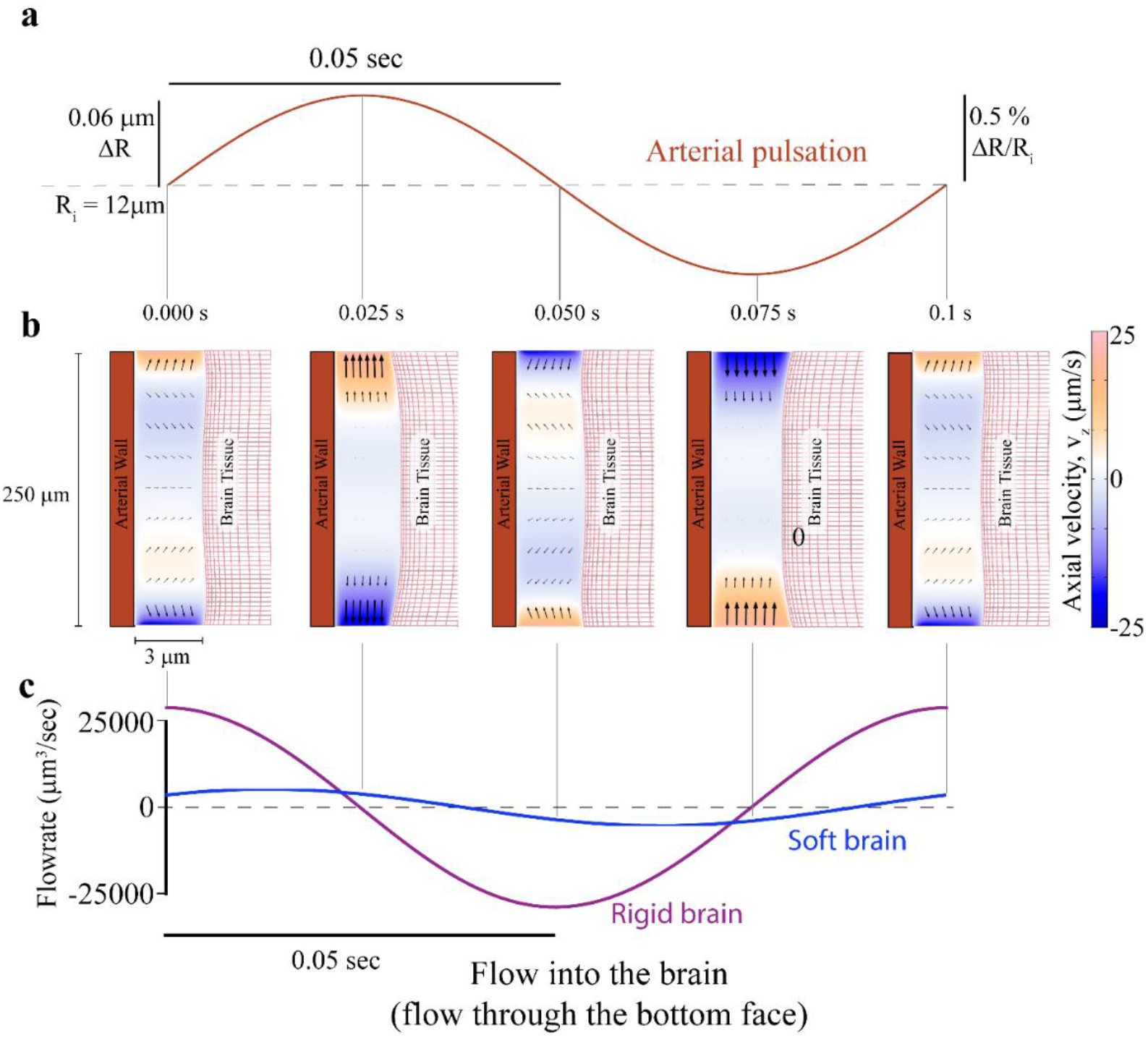
Minimal unidirectional perivascular fluid flow during arterial pulsation in a compliant brain model. This model uses a zero-pressure boundary condition on both ends of the PVS, to quantify the flow driven by peristalsis alone. Note the geometry is depicted with an unequal aspect ratio in the radial (r) and axial (z) directions for viewing convenience. **a.** Plot of the imposed heartbeat-driven pulsations in arterial radius (±0.5% of mean radius^1^,R_i_) at 10 Hz, the heartrate of an un-anesthetized mouse. The pulse wave travels at 1 meter per second along the arterial wall, into the brain^2,3^. **b.** Plot of the fluid velocity in the axial (z) direction induced in the PVS by arterial pulsation. Colors denote the axial velocity in a cross section through the PVS for one complete pulsation cycle. Fluid velocity vectors (arrows) are provided to help illustrate the flow direction. The fluid flow is nearly symmetric through both ends of the PVS throughout the cycle. The peak fluid velocity magnitude in this case is 6 times smaller than the one obtained when the brain is assumed to be rigid (Supplementary Fig 1b). Note: Arterial and brain tissue displacements induced by arterial pulsations are very small (<0.1 μm). To make the movements clearly visible, we scaled the displacements by 10 times in post-processing. **c.** Flow through the bottom surface of the PVS (into the brain) induced by the peristaltic movement of the arterial wall. The net flow over a single cycle was 0.03 μm^3^/sec. The peak instantaneous flow rates are an order of magnitude smaller than the values obtained in the rigid brain model (Supplementary Fig 1c).

**Figure S6.**
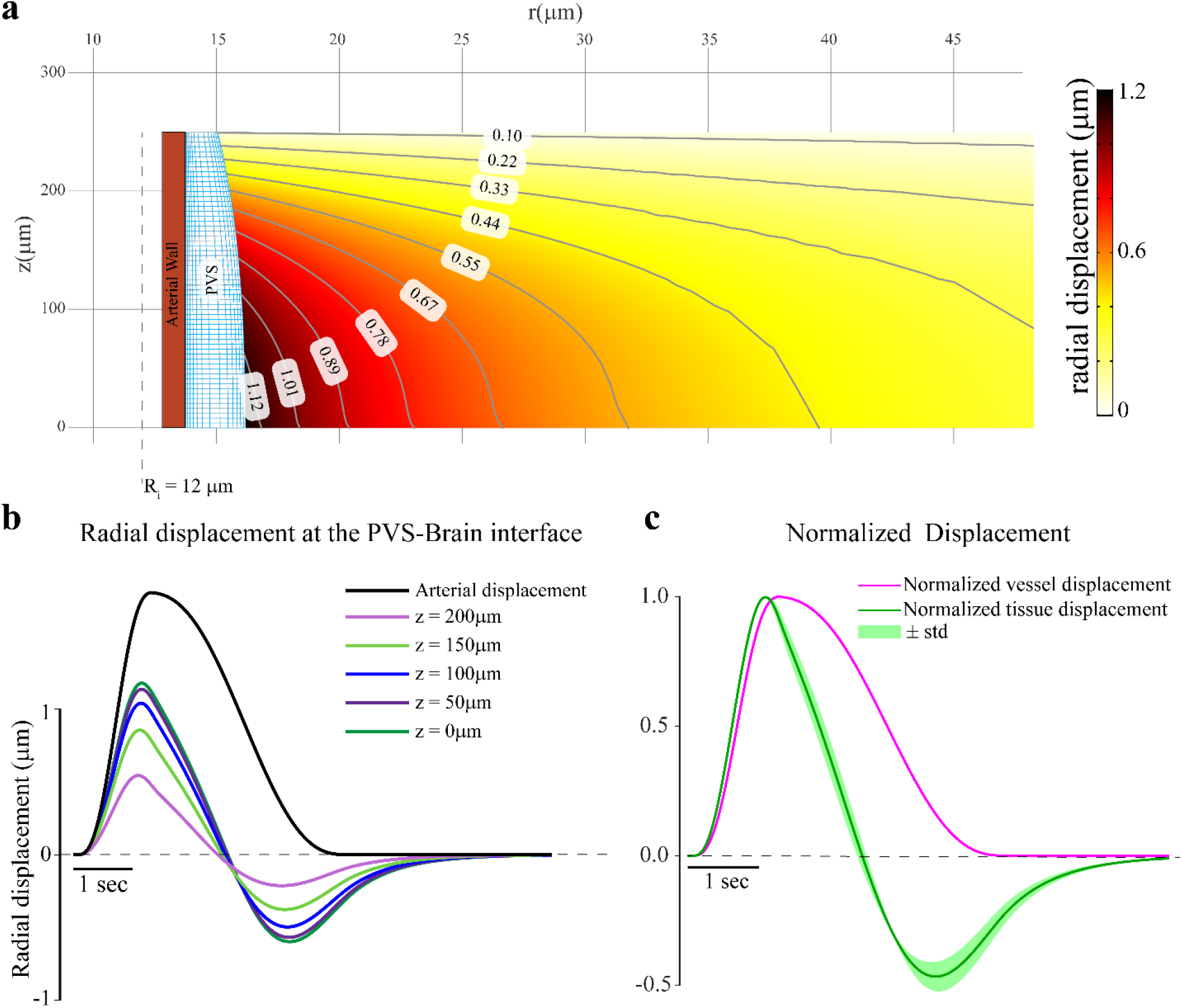
The pressure changes in the PVS due to fluid flow deform the brain tissue. Note the geometry is depicted with an unequal aspect ratio in the radial (r) and axial (z) directions for viewing convenience. a. Radial displacement contours in the brain tissue (maximum deformation, occurs at 1.16 seconds for the vasodilation profile shown in b). The brain tissue can deform by upto 1.2 μm, when the artery (with an initial radius of 12μm) increases its radius by by 1.8 μm. b. Plot shows the change of radial displacement at the PVS-Brain interface with time. These deformations can be explained by the pressure changes in the PVS. When there is fluid outflow from the PVS, the increase in the pressure causes the brain tissue to deforms radially outward and when there is fluid influx, the brain tissue deforms radially inward. c. Plot shows the mean and standard deviation of the normalized displacement taken at 100 different points in the brain tissue (10×10 grid between r = (15,65) μm and z = (25,225) μm in green. The normalized displacement of the arterial wall is shown in magenta. The displacement values are normalized by the peak displacement value at the location (L-infinity norm). This plot suggests that even though there is a large variation in the actual displacement of the brain tissue (shown in **a.** and **b.**), the normalized displacement response of the brain tissue is almost identical everywhere.

**Figure S7.**
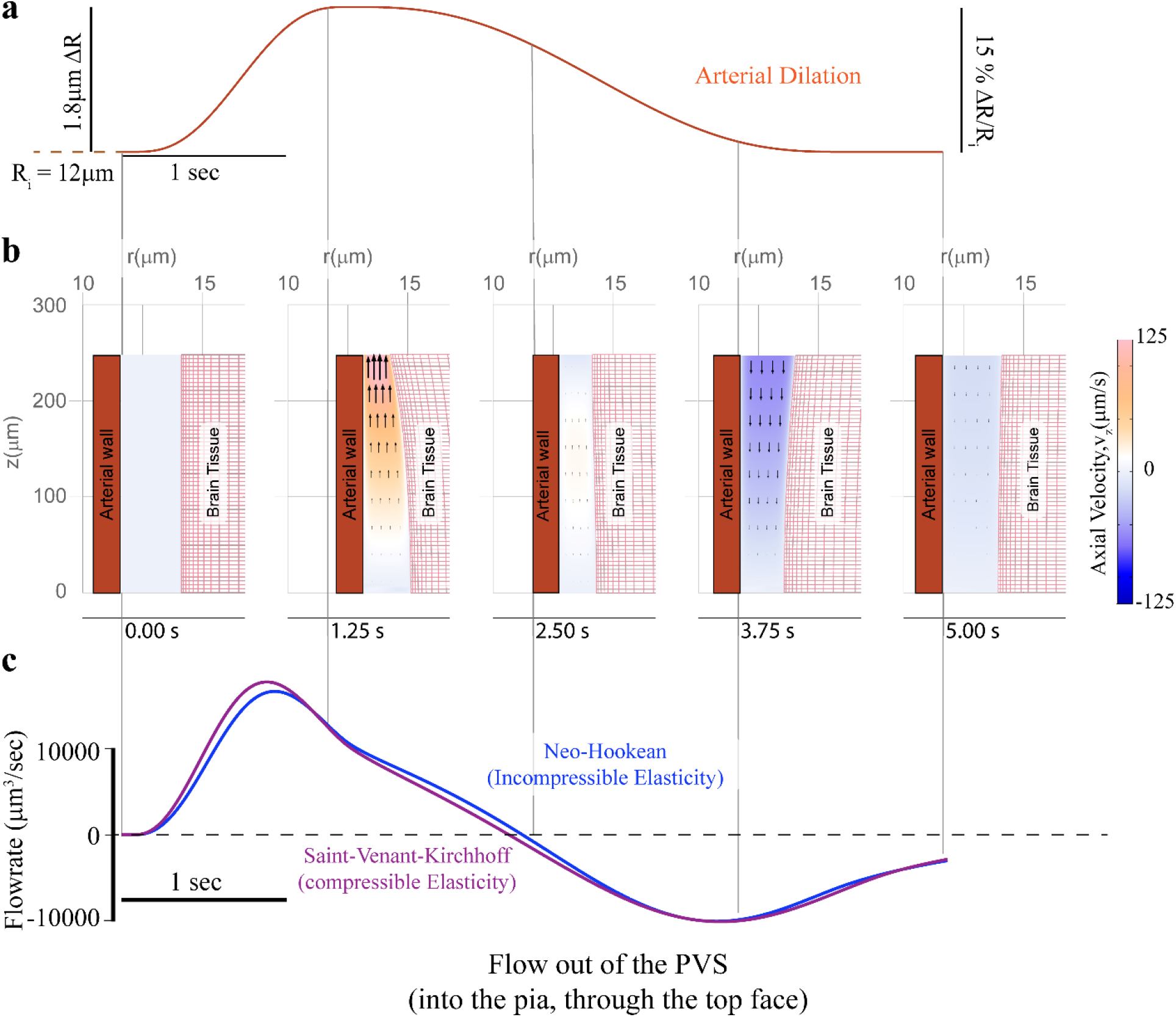
Vasodilation induced PVS fluid flow in a completely incompressible, Neo-Hookean model was very similar to the compressible SVK model. Note the geometry is depicted with an unequal aspect ratio in the radial (r) and axial (z) directions for viewing convenience. **a**. Plot of the proscribed arterial wall movement, which is identical to the one shown in Fig 4a. **b.** Plot showing the axial (z-direction) fluid velocity a cross section of the PVS, when the arterial wall movement is given by neural activity-driven vasodilation. A positive velocity value (orange) means that fluid leaves the PVS and negative value (blue) means that fluid enters the PVS. Fluid velocity vectors (arrows) are provided to help the reader interpret the flow direction from the colors. The region in white has little to no flow. These plots (very similar to the ones in Fig 4a) show that compared to heartbeat-driven pulsations (supp Fig 3b), vasodilation-driven fluid flow occurs through the entire length of the PVS and has substantially higher flow velocities. **c.** Flow out of the PVS and into the pia, through the top face of the PVS. The flow rates predicted by the model with nearly incompressible (SVK model with Poisson’s ratio of 0.45) brain tissue (magenta) and a completely incompressible, Neo-Hookean model (blue) are very similar.

**Figure S8.**
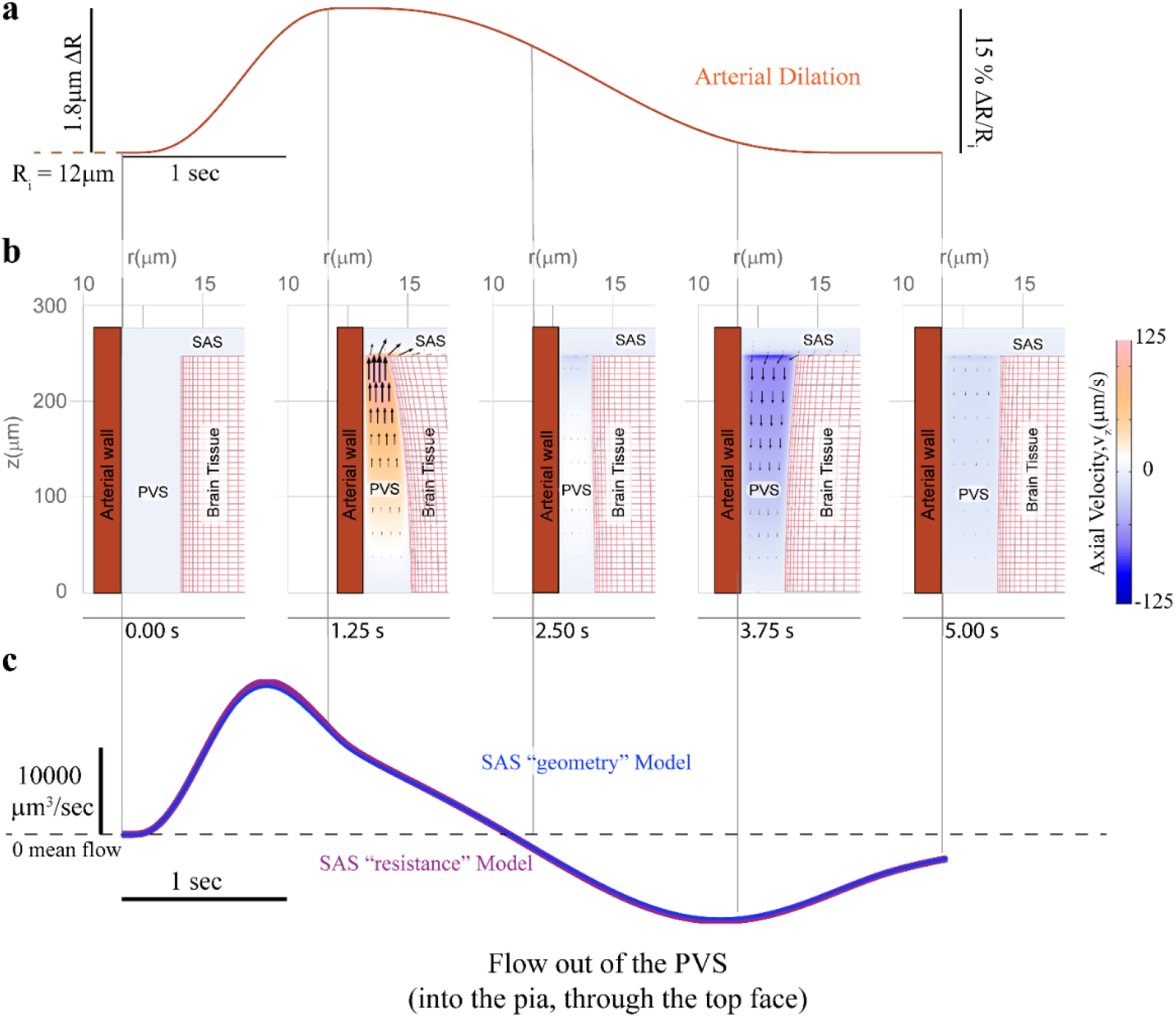
Arterial dilations during functional hyperemia drive fluid exchange between the PVS and SAS in the SAS “geometry” model. Note the geometry is depicted with an unequal aspect ratio in the radial (r) and axial (z) directions for viewing convenience. **a.** The arterial wall movement is prescribed by a typical neural activity-driven vasodilation response, the same one shown in Fig 4a. **b.** Plot showing the axial velocity (velocity in the z-direction) in a cross section of the PVS and the connected SAS, when the arterial wall movement is given by neural activity-driven vasodilation. A positive velocity value (orange) means that fluid leaves the PVS and negative value (blue) means that fluid enters the PVS. Fluid velocity vectors (arrows) are provided to help the reader interpret the flow direction from the colors. Because the fluid is incompressible, the flow speed decreases when flowing into the SAS, which has a larger area of cross section compared to the PVS. The region in white has little to no flow. These plots (very similar to the ones in Fig 4a) show that compared to heartbeat-driven pulsations (supp Fig 3b), vasodilation-driven fluid flow occurs through the entire length of the PVS and has substantially higher flow velocities. Note that the scale for the radial direction is different than that in the axial direction. **c.** Flow out of the PVS and into the pia, through the top face of the PVS. The flow rates predicted by the SAS “resistance” model (magenta) and the SAS “geometry” model (blue) are almost identical.

**Figure S9.**
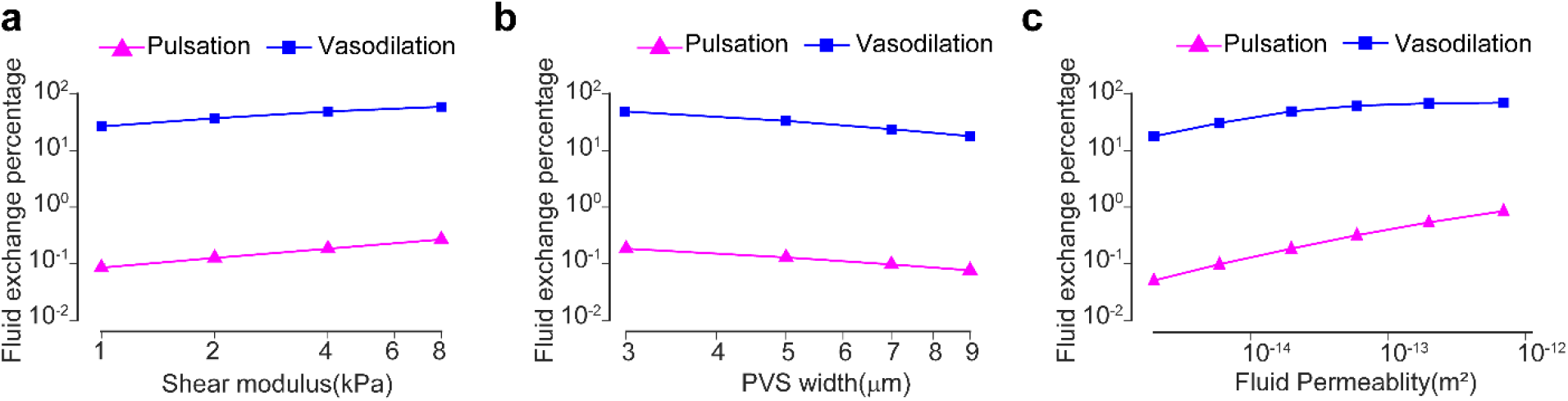
Vasodilation drives appreciably higher fluid exchange between the PVS and subarachnoid space compared to heartbeat driven pulsations. The plots show the changes in fluid exchange percentage, the percentage of fluid in the PVS exchanged with the SAS, with change of model parameters. The model predicts that compared to arterial pulsations, the vasodilation driven fluid exchange percentage is two orders of magnitude higher. This difference is similar for different values of elastic modulus of the brain(a), the width of the PVS (b) and the fluid permeability of the PVS (c).

**Figure S10.**
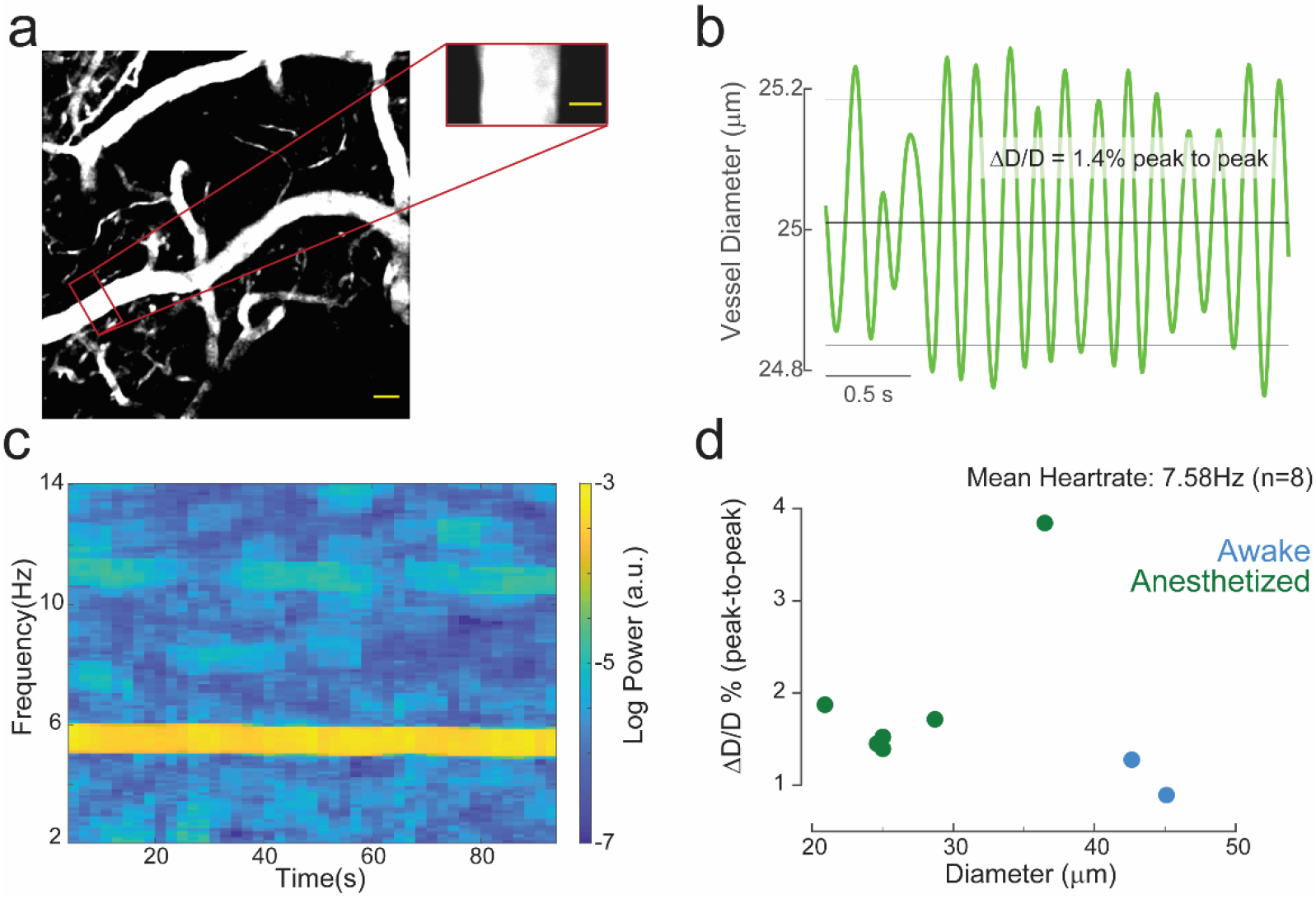
Heartbeat drives 0.5-4% (peak-to-peak) changes in arterial diameter. **a.** Sample image of the in-vivo fluorescence measured by two-photon microscopy following intravenous injection of FITC conjugated dextran (150 kDa) shows the cerebral vasculature near the surface of the brain (scale bar = 25 μm). Inset (scale bar = 10μm) shows a smaller region containing a segment of the artery, that is scanned at 30Hz to obtain arterial diameter changes in the typical heartrate frequencies (4-14 Hz). **b.** Sample plot of the diameter values measured for the artery shown in **a.** The plot shows that heartbeat drives 1.4% peak-to-peak change in diameter for this artery. **c.** Spectrogram shows the log power of diameter changes for the sample artery shown in a. There is a clear peak in spectral power at 5.59 Hz, which is the heartrate frequency. **d.** Scatter plot shows the relation between the percentage changes in diameter (8 vessels, 6 mice) and the mean diameter at heartrate frequencies. We found that the heartrate driven pulsations are smaller in awake animals(blue) compared to isoflurane-anesthetized animals(green). The pulsations in awake animals could only be measured in large arteries, probably because the pulsations in smaller vessels are below the image resolution attainable by our imaging equipment.

**Figure S11.**
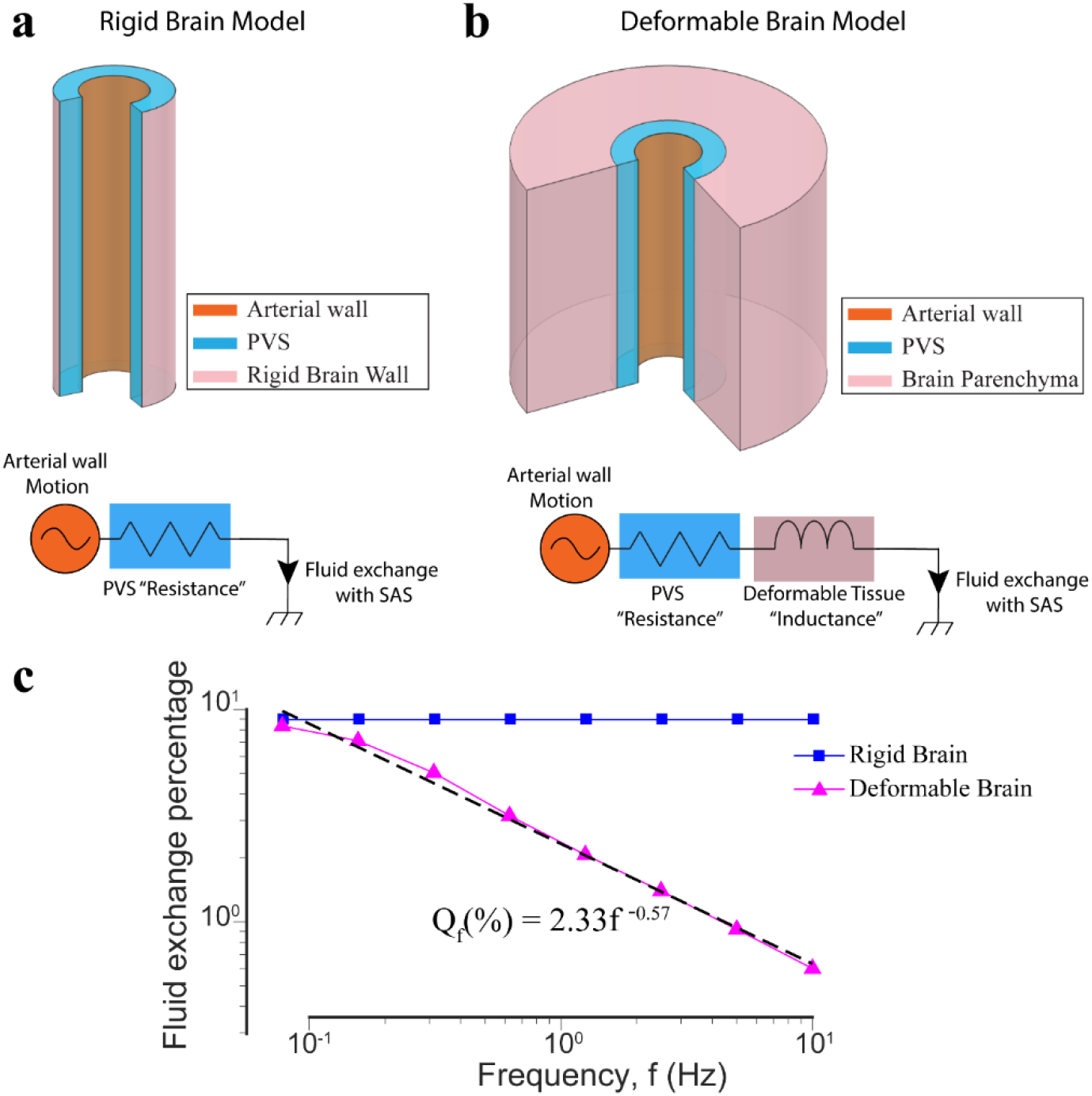
The presence of deformable brain tissue makes the PVS more resistant to fluid flow changing at high frequency. **a.** Geometry for a “rigid brain” model(top) and the equivalent circuit diagram. The driver for fluid flow is the arterial wall motion. The flow resistance of the PVS can be modelled by a simple resistor is independent of the frequency of the arterial wall movement. **b.** Geometry for the fluid-structure interaction model with a deformable brain(top) and the equivalent circuit diagram. The driver for fluid flow is the arterial wall motion. The total flow resistance of the system can be modelled by a resistance from the PVS and an inductance because of the deformable tissue. In this model, the flow resistance of the system increases with increase in the frequency of the arterial wall motion. This means that for arterial wall motion at high frequency, less fluid will be exchanged between the PVS and the SAS. **c.** Plot shows the relation between fluid exchange percentage and frequency of arterial wall motion. The arterial wall motion was given by a 4% peak-peak sinusoidal wave with different frequency values. The default values were used for all other parameters (see Table 1). For very low frequencies (<0.1 Hz), the fluid exchange driven by the arterial wall is same with a deformable brain tissue and a rigid brain. For higher frequencies, the fluid exchange percentage has an inverse power law relation with the frequency of arterial wall motion.

**Figure S12.**
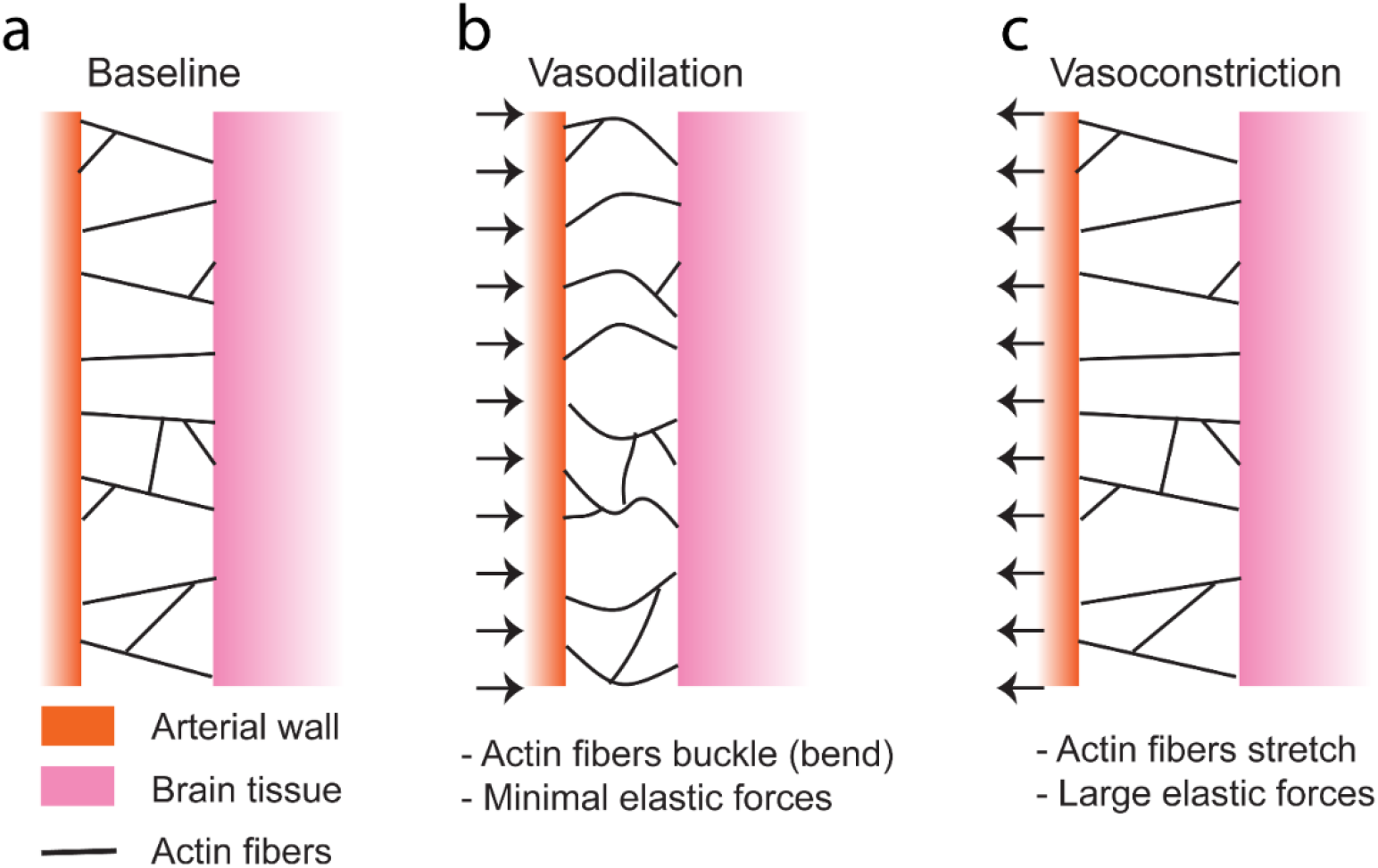
The lack of negative radial displacement in the brain tissue can be attributed to the non-linear elastic response of the connective tissue in the PVS. a. The connective tissue in the PVS is possibly made up of actin fibers. b. When arteries dilate, the connective tissue is under compression (middle) and the actin fibers buckle (bend) rather than compress due to the low energy cost of bending. Therefore, there are very low elastic forces and our assumption that the forces in the PVS originate mainly from the fluid pressure is valid. c. When the arteries constrict or return to their original size, the connective tissue is in tension and the actin fibers stretch, creating significantly larger elastic forces. In this case, our assumption that the forces in the PVS originate mainly from the fluid pressure does not hold and the fluid-structure interaction model cannot predict the behavior accurately.

**Figure S13.**
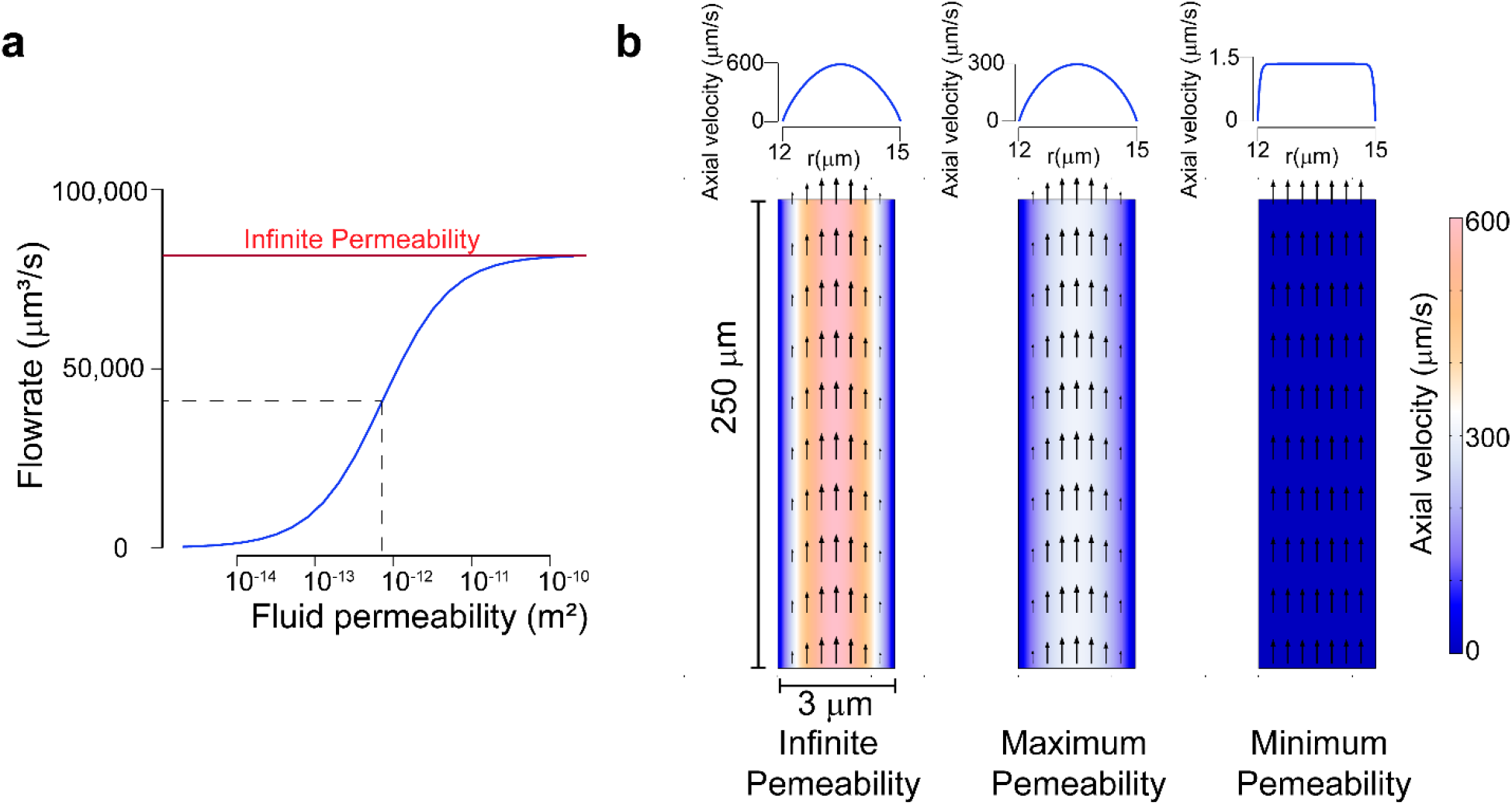
Choosing the maximum fluid permeability parameter for the PVS. We took the geometry of the PVS and applied a pressure difference of 1mm of Hg across the length. The fluid motion is governed by the Darcy-Brinkman equation with the fluid properties of CSF. Note the geometry is depicted with an unequal aspect ratio in the radial (r) and axial (z) directions for viewing convenience. **a.** Fluid flowrate through the top face of the PVS changes with the fluid permeability. The flow rate reaches its maximum possible value at infinite permeability. We chose the permeability that gives half the flowrate at infinite permeability (7×10-13 m^2^) as the maximum possible fluid permeability of the PVS. **b.** Plots show axial velocity color plots for a pressure difference of 1 mm of Hg across the ends of the PVS. The three plots are for infinite permeability, maximum (7.0×10^−13^ m^2^) and minimum (2.0×10^−15^ m^2^) permeability used in the simulations in this article. Fluid velocity vectors (arrows) are provided to help the reader interpret the flow direction from the colors. The line plots on the top show the axial velocity profile for the corresponding permeability values.

**Figure S14.**
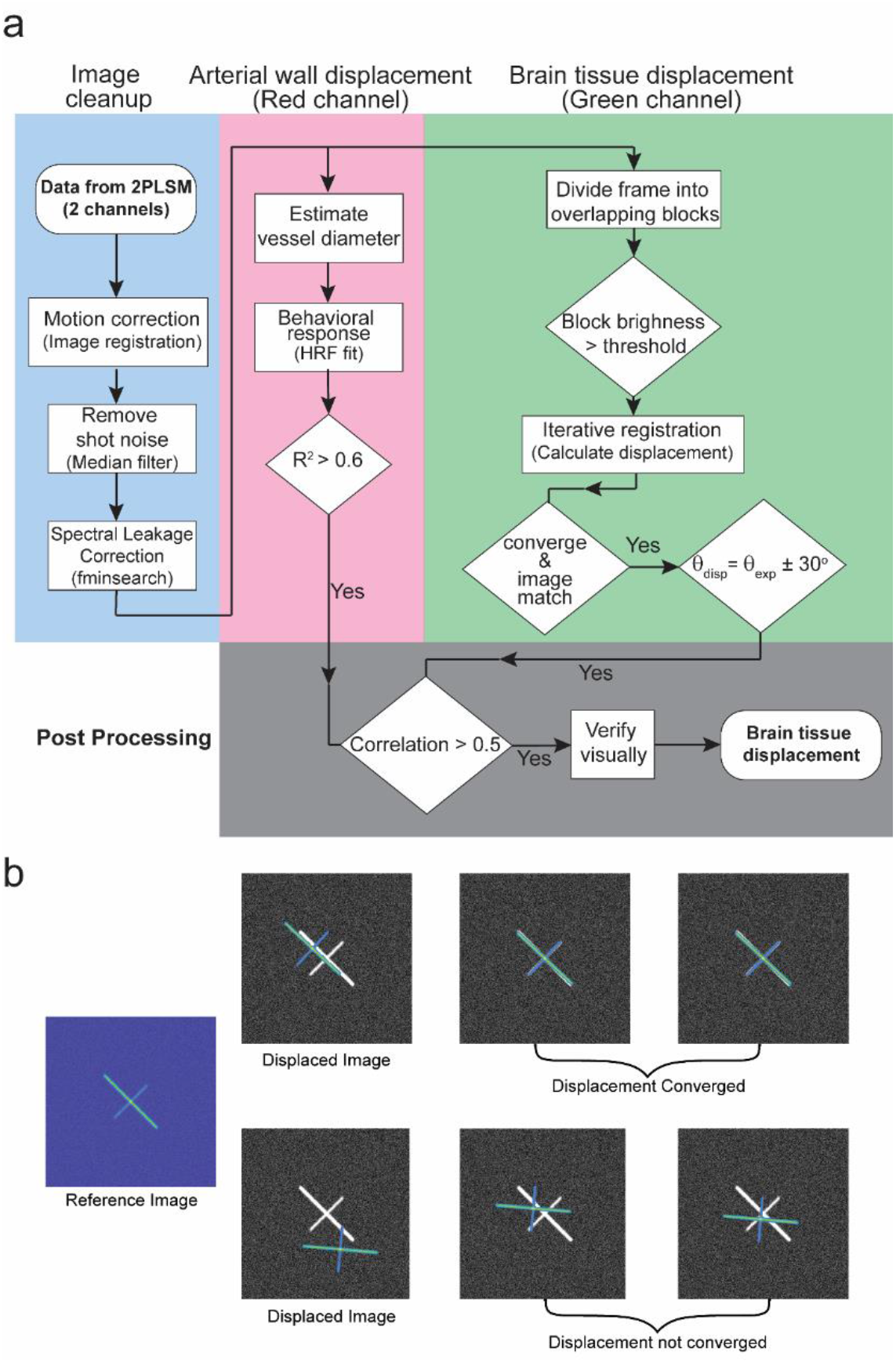
The procedure for measuring brain tissue displacement from in-vivo imaging data collected with a two-photon laser scanning microscope. **a.** Flow chart depicting the complete procedure used to calculate displacements in the brain tissue. The procedure can be broken down into 4 major sub-sections as shown in the figure. For a full description of the procedure, see methods. **b.** A depiction of the iterative method in calculating displacements. The figure on the left shows a reference image. The intensity is shown by a Parula colormap (Matlab). The images on the right show two cases of displaced images. The one on the top is rotated by 2°, and can be matched to the reference image (shown in gray) by a simple displacement. After the first calculation of the displacement and correcting the displaced image, the reference and the displaced image match and further iterations of displacement calculation yield a zero value, showing that the displacement calculation has converged. The one on the bottom is rotated by 45°, and cannot be matched to the reference image (shown in gray) by a simple displacement. In this case, every iteration of displacement calculation yields a non-zero value and the calculation is not converged.

**Figure S15.**
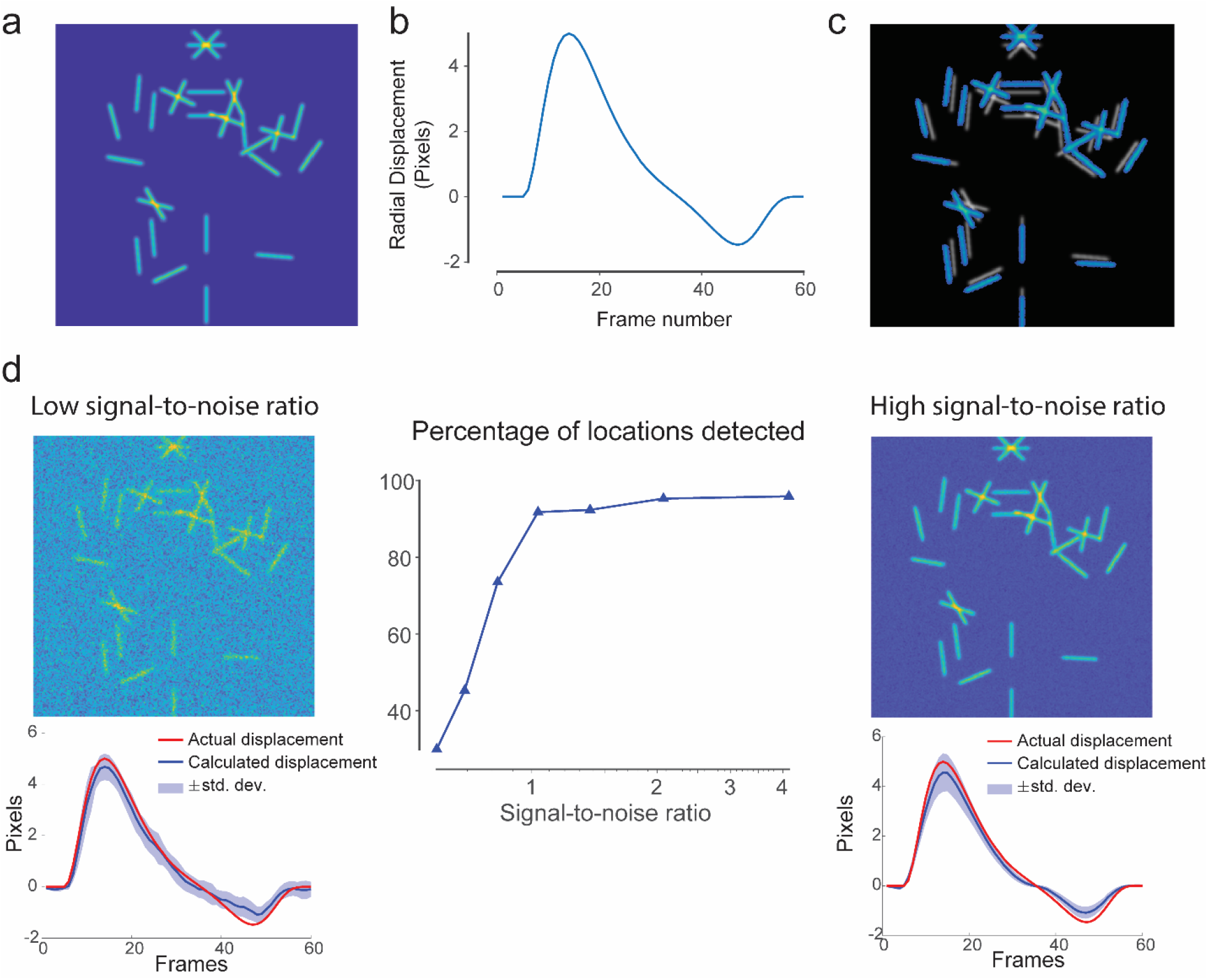
Applying the displacement calculation procedure on pseudo data shows that the method is robust to noise. **a.** A computer generated image (512×512 pixels) with randomly oriented lines. **b.** The radially-outward displacement given to the image shown in **a.** **c.** An image showing the radially-outward displacement at peak displacement (frame number 13). The initial position of the lines is shown in white and the displaced position is shown in blue. **d.** The displacement extraction procedure (shown in Supplementary figure 12) is robust to noise and predicts correct displacement. On the left, a case with low signal-to-noise ratio (0.59) is shown. The calculated displacements are very close to the actual displacement. The accuracy is comparable to the case with high signal-to-noise ratio (4.14) on the right. However, high noise results in a detection of displacement at fewer locations. The plot in the center shows that at low signal to noise ratio only 30% of the possible locations can be used for displacement calculations. Signal-to-noise ratio is calculated as the ratio of the mean signal value to the standard deviation in the noise.

## Appendix

## A Kinematics in ALE

To describe the motion of a continuum(solid, fluid), we start with two configurations, the deformed configuration *B_t_*, the reference configuration *B_p_*. As shown in Fig 1, the deformed configuration *B_t_* is the image of *B_p_* under the map ***χ**_p_*. The points in *B_p_,B_t_* are denoted by ***X**_p_, **x*** respectively. The displacement of material particles from the reference to the deformed configuration is given by.

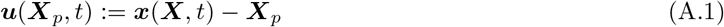

Alternatively, the Eulerian description of the displacement is given by

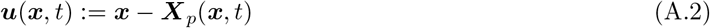

Where ***X**_p_*(*x, t*) is the image of *x* under the map 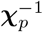 at time t.

Denoting the material time derivative by *D_t_*, the material particle velocity ***υ*** is related to the Eulerian description of the displacement field as follows

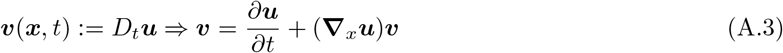

where **∇**_*x*_ is the gradient with respect to *x* over *B_t_*. The above equation for the material time derivative holds true for the Eulerian description of any vector quantity. It can be viewed as a property of the map ***χ**_p_*

Additionally, for the ALE computation, we have the computational configuration *B_c_*. The points in *B_c_* are denoted by ***X**_c_*. As shown in Fig. A.1, the deformed configuration *B_t_* is the image of *B_c_* under the map *χ_c_*. In a finite element computation, the map *χ_c_* can be interpreted as the motion of the mesh. The displacement of the mesh gives us the relation between position of a particle in the computational domain and its position in the deformed configuration.

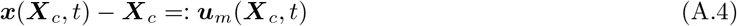

The deformation gradient and the Jacobian determinant of the mesh motion are given by

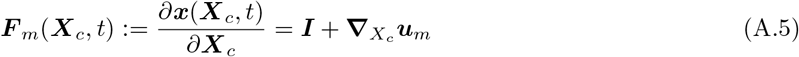

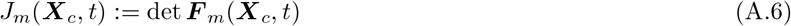

where, **∇**_*X_c_*_ is the gradient with respect to ***X**_c_* over *B_c_*.

Now, we take the material time derivative of all the quantities in eq. A.4. The material time derivative of *x*(***X**_c_,t*) is the ALE representation of material particle velocity.

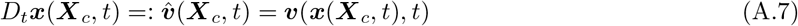

**Figure A.1:**
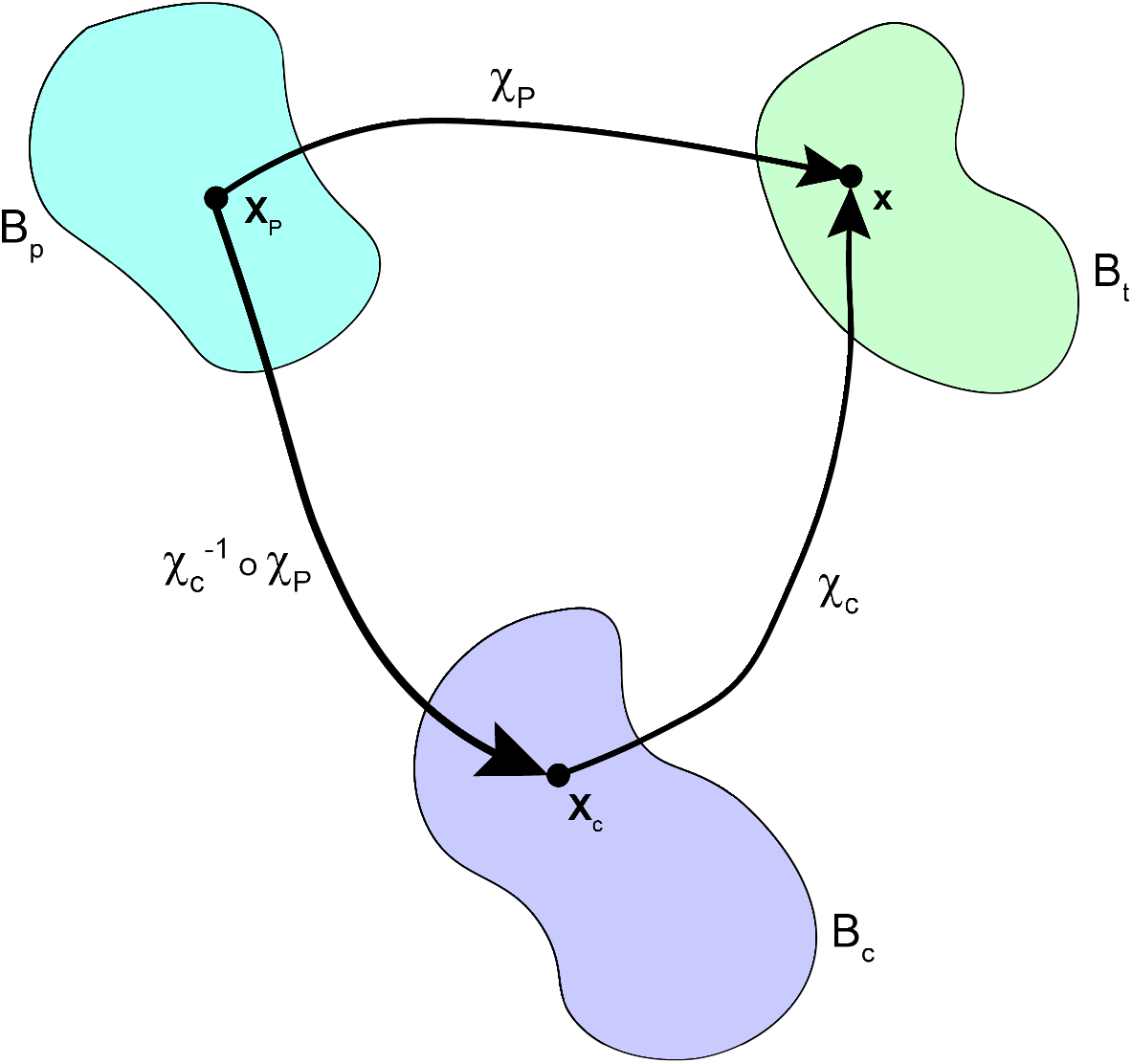
Current configuration *B_t_* is the common image of particle motion *χ_p_* and mesh motion *χ_c_*

We want to find the solution to an initial boundary value problem(or a boundary value problem) on the deforming domain. However, since our finite element computation is performed on the computational domain *B_c_* instead of the deforming domain *B_t_*, the primary unknown fields,for example, the stress tensor (***σ***) fluid velocity (***υ***) and pressure (*p*) will be calculated as functions of the computational coordinates ***X**_c_* instead of the physical coordinates ***x***, and time *t*.

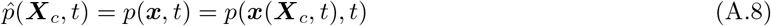

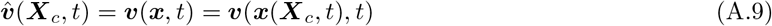

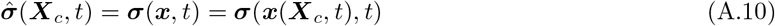

We will need to express the governing equations in continuum mechanics, that contain spatial gradients and material time derivatives, in ALE coordinates. The following transformations use the deformation gradient defined in eq. A.5 and the Jacobian determinant defined in eq. A.6 to rewrite the spatial derivatives(that commonly occur in mass and momentum balance laws) in the computational domain.

For a vector ***υ***:

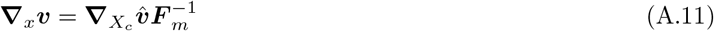

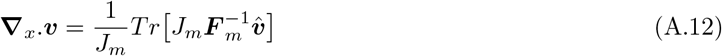

where, Tr is the trace operator on tensors.

For a scalar *p*:

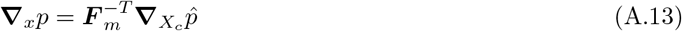

For a tensor ***σ***

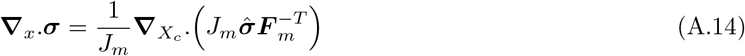

The material time derivative of velocity(the acceleration) estimated in the physical and computational coordinates is given as follows.

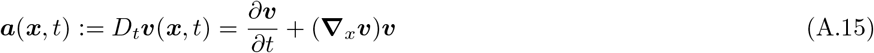

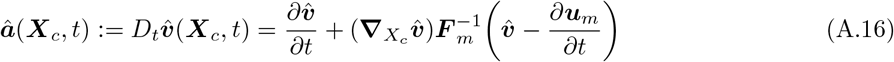

The derivation of eq. A.16 is not immediately obvious. A detailed derivation is given by Donea *et al*. [3].

Further, any integral over the deformed domain can be transformed to an integral over the computational domain using the Jacobian determinant defined in eq. A.6.

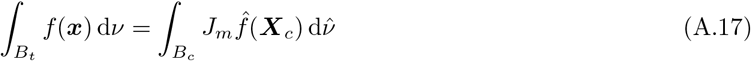

where *ν* and 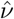 are infinitesimal volumes is *B_t_* and *B_c_* respectively.

An integral over the boundary of the deformed domain can be transformed into an integral over the boundary of the computational domain as follows

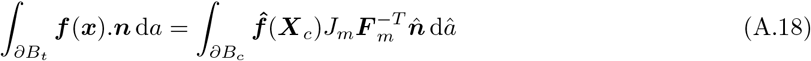

where *a* and *â* are infinitesimal areas in *∂B_t_* and *∂B_c_* respectively and 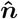 is a unit normal in the computational domain.

## B Particle tracking in ALE

To track fluid particles in the computational domain(in ALE), we want to find the fluid particle coordinates (***X**_c_*) as a function of time. The material time derivative of ***X**_c_* is the particle velocity observed from the computational domain *B_c_* is defined as:

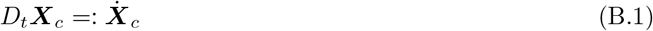

The material time derivative of the mesh displacement is obtained by a method similar to that in eq. A.3. In this case we use the map 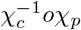 instead of the map *χ_p_* in eq.A.3.

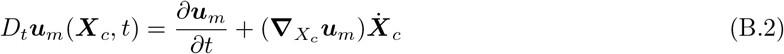

From the finite element computation in ALE, we know the fields 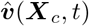 and ***u**_m_*(***X**_c_,t*). We want to find 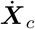 as a function of these known quantities. Using eq.s A.7, B.1 and B.2 in the material time derivative of eq. A.4 and rearranging the terms, we get

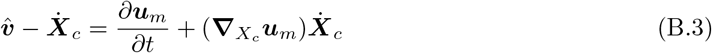

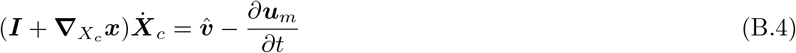

We can use the definition of ***F**_m_* from eq. A.5 to obtain the particle velocity as seen from the computational domain

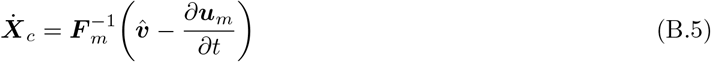

Here, we present the initial value problem to calculate fluid particle trajectories in ALE framework. This calculation is proposed as a post-processing step after the actual finite element computation. Therefore, it is assumed that the material particle velocity 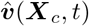 and the mesh displacement ***u**_m_*(***X**_c_, t*) (and subsequently ***F**_m_* and 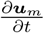) are known quantities in the computational domain.

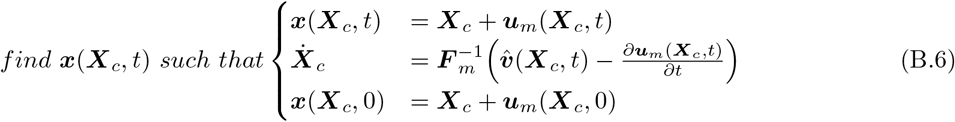

The above initial value problem can be solved using a forward Euler integration scheme to obtain fluid particle position in the computational and deformed domains. The time integration scheme is implemented as follows to calculate the position in the computational domain at time t+dt when the position at time t is known.

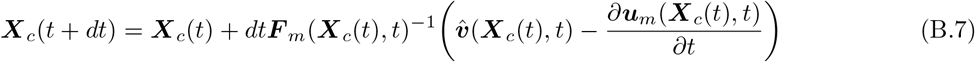

The particle position in the deformed domain is simply a consequence of the first equation of the initial value problem(eq. B.6). We can track fluid particles till the time they exit the computational domain.

There are two main reasons for choosing a forward Euler integration scheme over a backward Euler integration scheme. Firstly, the ODE described in eq. B.6 does not necessarily have a fixed point. Secondly, the values of the quantities on the right side of the ODE only exist within the computational domain, and therefore the equation is not solvable using a backward Euler scheme at the timestep when a particle leaves the computational domain.

## C Initial conditions for problems

In all the intial-boundary value problems described below, the initial values and initial time-derivatives of all variables are always set to zero. This is chosen for convenience since specifying a non-zero initial condition for one variable would require the knowledge the corresponding initial values for all the other variables with coupled physics. The parameter values used in Dirichlet boundary conditions, are ramped up from zero to the specified values using step functions in Comsol multiphysics. These functions have continuous first and second derivatives with respect to time. In the subsequent sections, we will use *step*1(*t*) in the equations to indicate wherever these functions are used. The results for these simulations are shown after the effect of the initial conditions are insignificant. We show the results after 20 cycles of the peristaltic wave, where the variations in the velocity and pressure fields from cycle to cycle are less than 10^−6^% of peak value.

## D Darcy-Brinkman Flow in ALE coordinates

We want to solve for the fluid velocity *ν_f_* and pressure *p_f_* in a deforming domain *B_t_*, representing the PVS. The displacement field that defines the time-dependent deformation of the domain is denoted by ***u**_m_* (same as eq. A.4). The finite element calculations are done in the computational domain *B_c_*, where we solve for the velocity and pressure fields as a function of the computational coordinates ***X**_c_*. These fields 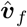 and 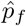 are defined according to equations A.9 and A.8 respectively.

The computations are done in an axisymmetric framework. The inner and outer radius of the computational domain are *R*_1_ & *R*_2_ respectively, where *R*_2_ = *R*_1_ + *wd*. The domain has a length *L_a_* in the *z* direction. See Table 1 for the list of parameters

We use a harmonic model for the mesh motion [1, 2].The governing equation for the mesh displacement, ***u**_m_* on *B_c_* is given by

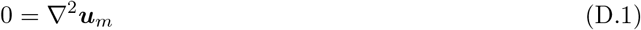

From here on in the document, since all the equations are presented in the computational coordinates, the *X_c_* subscript on the spatial derivatives (gradient, divergence and laplacian) is dropped.

The boundary of *B_c_* (*∂B_c_* =: Γ_*c*_) is divided into four non-overlapping regions such that 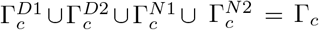, and 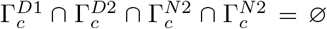, where, *∅* is the empty set. The division of the boundary is shown in Fig D.1. The mesh displacement on the boundaries should reflect the motion of the physical domain. Here, the outer wall of the artery 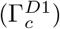 moves according to the sinusoidal wave described by eqn D.2.

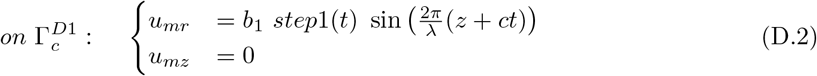

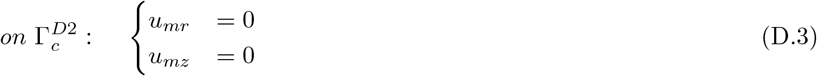

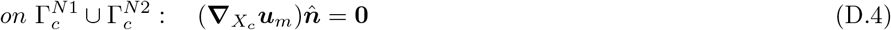

**Figure D.1:**
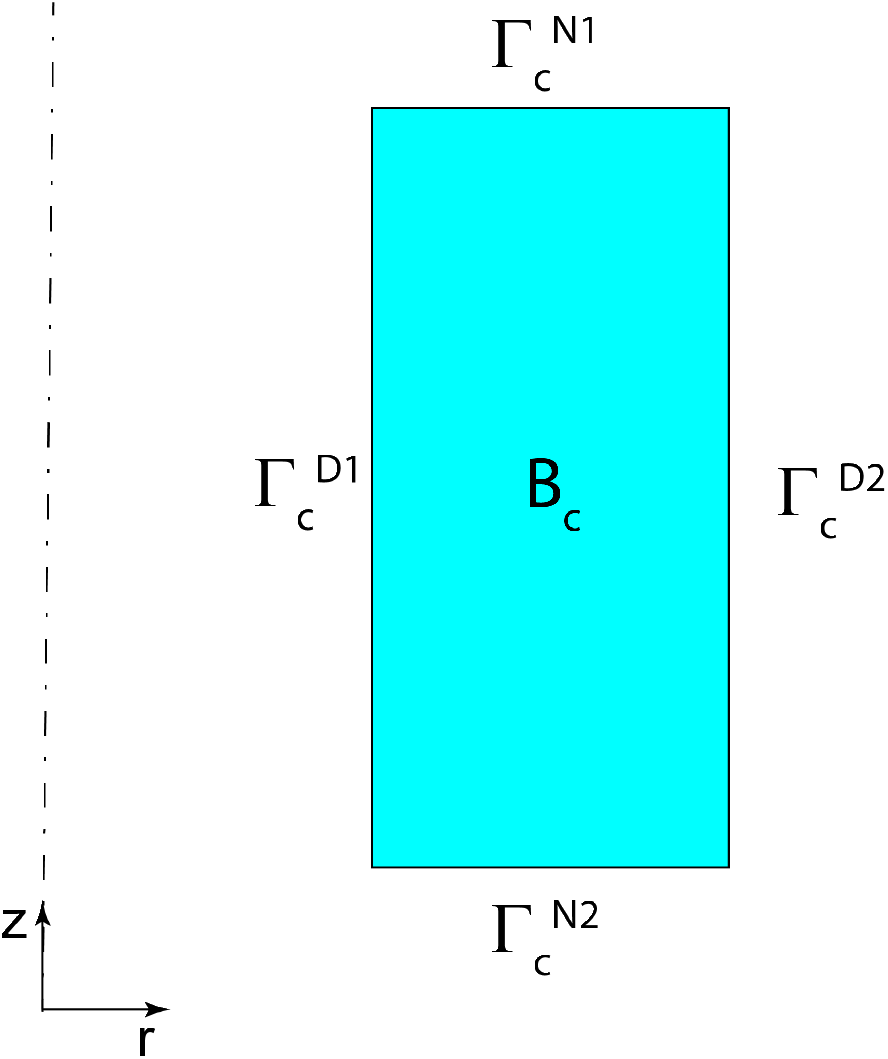
Computational Domain and boundaries for the Darcy-Brinkman flow problem ALE

The governing equation for the velocity 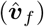 and pressure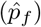 fields are given by **incompressible Darcy-Brinkman’s flow**. The Eulerian form of the governing equations for incompressible Darcy-Brinkman flow are well known.

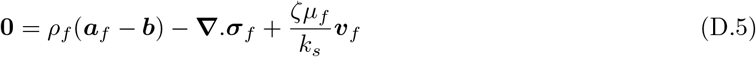

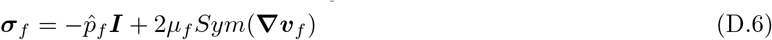

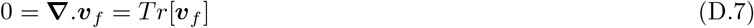

By converting all the derivatives to the computational coordinates(eqs A.11–A.18), we get the ALE formulation.

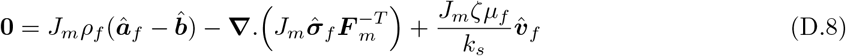

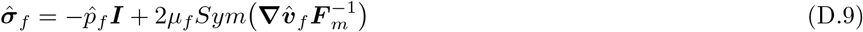

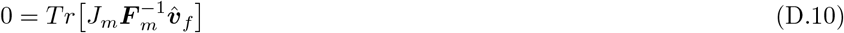

Where ***â**_f_* is the acceleration of fluid particles defined according to eq. A.16 and ***I*** is the identity tensor. The body force, ***b*** is zero for our problem.

We use a no-slip boundary condition at the arterial wall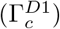 and the brain tissue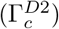 boundaries of the PVS. At the axial ends of the PVS, we assume that there is no applied traction.

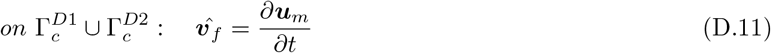

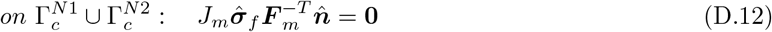

The weak form of equations D.1, D.8 and D.10 are solved with the boundary conditions in equations D.2–D.4, D.11 and D.12. We discretize the 2D(axisymmetric) geometry using Lagrange polynomials of second order for ***u**_m_* and 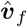, and Lagrange polynomials of first order for 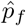. We use a fully coupled time dependent solver and a backward difference time integration scheme with a time step of 0.001s.

## E Fluid Structure Interaction

We have two domains *B_f_* and *B_s_*, representing the PVS and the Brain tissue respectively. We want to solve for the velocity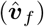 and pressure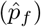 fields in *B_f_*. We continue to use the hat notation for these fields because they are calculated in the computational domain that represents the un-deformed PVS. We use the Darcy-Brinkman flow model for the fluid dynamics. The displacement of the computational domain is given the field ***u**_m_*. In *B_s_*, we want to calculate the solid displacement ***u**_s_* and solid velocity *υ_s_*. Saint-Venant-Kirchoff model is used for the solid elasticity.

For this problem, we use 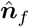 and 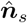 to represent the outward normals to the boundaries of *B_f_* and *B_f_* respectively.

**Figure E.1:**
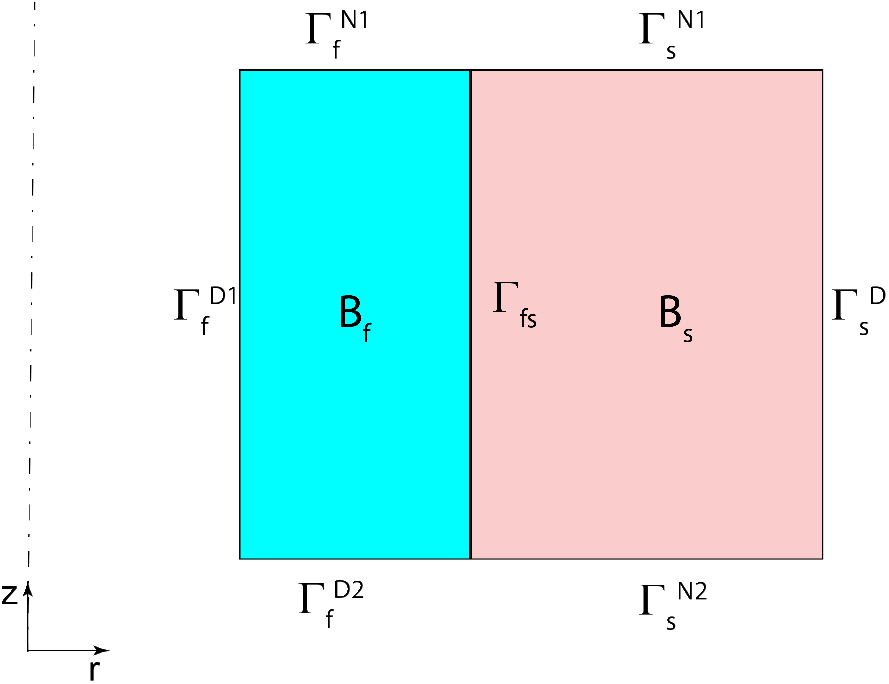
Computational Domains and boundaries for the Fluid-Structure interaction problem in section 4.1.4-4.1.6

The governing equations for 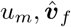 and 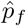 are given by equations D.1, and D.8–D.10. The boundary conditions at the fluid-structure interface (Γ_*fs*_) will be discussed at the end of this section. The boundary conditions on the remaining boundaries of *B_f_* are

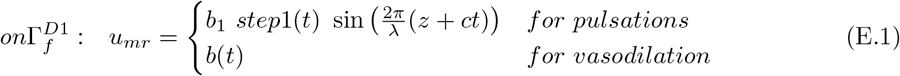

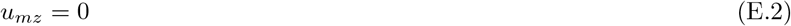

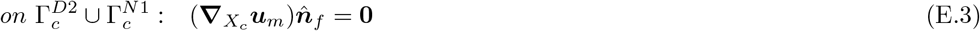

The function *b*(*t*) for vasodilation starts with zero initial value and is shown in main Fig 4.

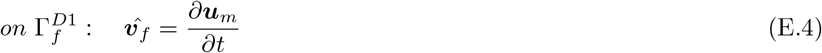

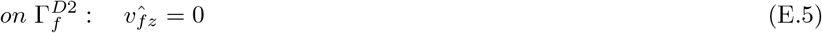

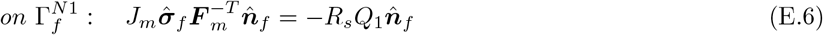

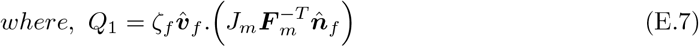

Here, *Q*_1_ is the flow rate out of the PVS through the face 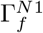. *R_s_* is the resistance of the subarachnoid space, which is taken to be 1% the flow resistance of the PVS. Equation E.6 is a Robin boundary condition, where the traction at a surface is proportional to the flow rate out of the surface. It serves as a lumped model for the subarachnoid space.

For the domain *B_s_*, the mesh motion is given by the solid displacement ***u**_s_*. This is the Lagrangian framework, used commonly in solid mechanics. The deformation gradient ***F**_s_* and the Jacobian determinant *J_s_* are defined similar to those in eqs A.5 and A.6. The strain energy for the Saint-venant-Kirchoff elastic model and the first Piola-Kirchoff stress are given by

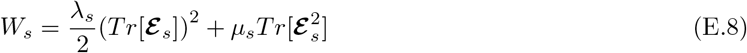

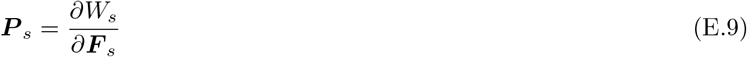

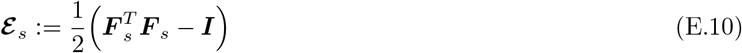

here, *Tr* is the trace operator in tensors.

Since the mesh in the domain *B_s_* follows the material particles, the material time derivatives in *B_s_* are equal to the partial derivatives. The governing equations for ***u**_s_, **v**_s_* are given by,

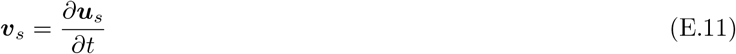

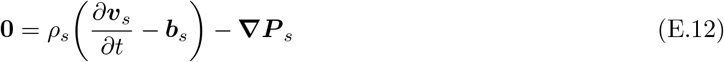

For our problem, the body forces *b_s_* are zero in *B_s_*.

The boundary conditions on Γ_*s*_ – Γ_*fs*_ are as follows.

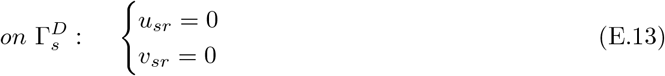

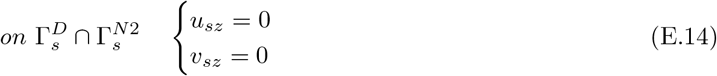

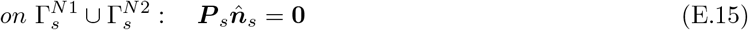

At the solid fluid interface, we enforce continuity of velocity and traction.

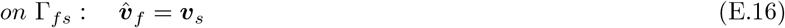

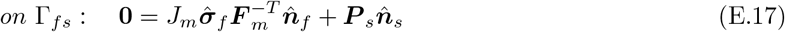

The weak form of equations D.1, D.8, D.10 on *B_f_* and E.11 and E.12 on *B_s_* are solved with the boundary conditions in equations E.1–E.7 and E.13–E.17. We discretize the 2D(axisymmetric) geometry using Lagrange polynomials of second order for 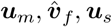 *and **υ**_s_* and Lagrange polynomials of first order for 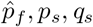 *and j_s_*. We use a fully coupled time dependent solver and a backward difference time integration scheme with a time step of 0.0001s for arterial pulsations and 0.0001s for functional hyperemia.

## F Volume exchange fraction

We are interested in calculating the maximum fraction of the fluid in the PVS exchanged with the SAS. To do this, we calculate the change of the fluid volume in the PVS at every time step, divide it by the initial volume of fluid in the PVS, and find the maximum value of this ratio.

Initial volume of fluid in the PVS can be calculated in the undeformed configuration of the fluid domain *B_f_*. For an fluid volume fraction(porosity) of *ζ*, we have the initial volume of fluid *V_i_*

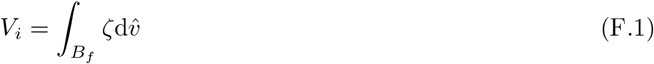

Since only the fluid leaves the PVS, the change in the fluid volume of the PVS is equal to the change in the total volume of the PVS. The calculation of the initial and deformed volume of the PVS is straight forward from A.17. The change in the PVS volume, *δV_t_* at any time t can be calculated over the undeformed configuration of the fluid domain *B_f_*.

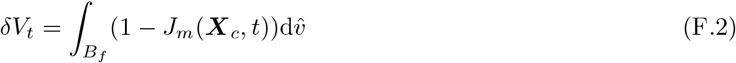

From these two quantities, we can calculate the fraction of PVS fluid volume exchanged with the SAS at any time t. The volume exchange fraction *Q_f_* is the maximum value of this quantity over the entire time period of the simulation (0 < *t* < *T*).

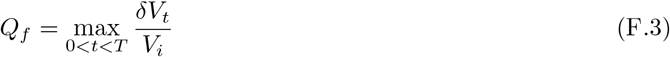

